# Binary outcomes of enhancer activity underlie stable random monoallelic expression

**DOI:** 10.1101/2021.08.27.457979

**Authors:** Djem U. Kissiov, Alexander Ethell, Sean Chen, Natalie K. Wolf, Chenyu Zhang, Susanna M. Dang, Yeara Jo, Katrine N. Madsen, Ishan D. Paranjpe, Angus Y. Lee, Bryan Chim, Stefan A. Muljo, David H. Raulet

**Affiliations:** Division of Immunology and Pathogenesis, Department of Molecular and Cell Biology, University of California, Berkeley, Berkeley, CA 94720, USA; Cancer Research Laboratory, University of California, Berkeley, Berkeley, CA, 94720, USA; Laboratory of Immune System Biology, National Institute of Allergy and Infectious Diseases, National Institutes of Health, Bethesda, Maryland 20892, USA

## Abstract

Mitotically stable random monoallelic gene expression (RME) is documented for a small percentage of autosomal genes. Here we investigated the role of enhancers in the RME of natural killer (NK) cell receptor genes. Enhancers were accessible and enriched in H3K27ac on silent and active alleles alike, decoupling enhancer activation and expression. Enhancers controlled gene expression frequency, as predicted by the binary model of enhancer action, and enhancer deletion converted the broadly expressed *Nkg2d* into an RME gene, recapitulating natural variegation. The results suggested that RME is a consequence of general enhancer properties and therefore many genes may be subject to some degree of RME, which was borne out by analysis of a panel of genes previously thought to be universally expressed within defined hematopoietic lineages: *Nkg2d*, *Cd45, Cd8a* and *Thy1*. We propose that previously documented RME is an extreme on a continuum of intrinsically probabilistic gene expression.

## Introduction

In most cases both alleles of autosomal genes are co-expressed. In recent years random monoallelic expression (RME) has emerged as an important exception that may apply to ∼0.5-10% of expressed genes in a given tissue and has been characterized as the autosomal analog of X-inactivation (*1*). In RME, different cells of a given type express only one allele, both or neither, and this expression pattern is mitotically stable. Notably, RME genes do not share an overarching unifying feature or function (*1–6*), and the biological role or RME is in most cases not known. RME is distinct from X-inactivation, genomic imprinting, and allelic exclusion of antigen receptor and odorant receptor genes, in that biallelic expression occurs at an appreciable frequency, and expression is largely stochastic rather than imposed by strict feedback regulatory mechanisms (*1*).

The molecular determinants of RME are poorly understood, in part because of difficulties analyzing single primary cells (*7*). Chromatin analysis of RME alleles in primary populations has not been possible due to the difficulty of isolating pure cell populations *ex vivo* with defined RME expression patterns (*1, 4*). As a result, previous analyses have been limited to clonal cell lines derived from F_1_ hybrids, where allelic expression is known and clonally stable (*4, 8–10*). These analyses revealed that enhancers of RME genes are constitutively accessible irrespective of gene or allelic expression status, whereas promoters are accessible only at active alleles (*4, 10*). Therefore promoter accessibility, rather than enhancer opening and activation, might be the “gatekeeper” of RME, whereas enhancers, being constitutively open, were proposed to permit rather than impose expression of RME alleles (*4*).

Enhancers may play more than a permissive role in RME, however, in light of evidence that enhancers primarily influence the probability of mitotically stable expression, rather than the amount of expression per cell (*11, 12*). In fact, deletions of enhancers resulted in mitotically stable gene variegation in both cell lines and normal tissues—notably *Igh* in B cell hybridomas and *Cd8a* in primary thymocytes, among others (*11–17*). Collectively, these data support the binary or “on/off” model of enhancer action (*18, 19*), where an increase in enhancer activity at a genetic locus results in an increase in the *probability* of gene expression, rather than an increase in expression per cell. Conversely, weak or reduced enhancer activity results in a lower likelihood of expression, but the cells that express the gene express a similar amount of gene product.

We reasoned that the binary action of enhancers—when limiting—might be a driving principle of RME, and sought to test this in an example of RME with a clear biological purpose: the *Ly49* and *Nkg2a* receptor genes expressed by NK cells (*1, 6, 20*). These genes, clustered in a ∼1 Mb stretch of the NK gene complex (NKC) on chromosome 6 in mice, encode cell surface receptors that bind MHC I molecules. They are expressed in a variegated (*21, 22*), monoallelic (*23*), stochastic and largely mitotically stable fashion (*24*), resulting in subpopulations of NK cells that express random combinations of the receptors and consequently exhibit distinct reactivities for cells expressing different MHC I molecules. Regulation of each gene allele is independent, and expression of one *Ly49* gene has minimal effects on expression of others (*25*). While a clear biological purpose for RME at many genes is lacking, RME at *Ly49* genes underlies the basis of the “missing self” mode of NK cell target recognition (*26*). Furthermore, the system represents a powerful *in vivo* genetic model of RME, where allelic expression states can be easily assessed at the population level in primary cells using allele-specific antibodies that we and others previously generated (*25, 27*), circumventing previous technical limitations to studying RME in single primary cells.

Importantly, competition between Ly49 genes for interaction with a shared enhancer or locus control region is not required for variegation of *Ly49* genes, as a *Ly49a* genomic transgene ectopically integrated in different genomic sites was usually expressed with a frequency similar to that of the native *Ly49a* gene (∼17% of NK cells) (*28*). We previously identified a key DNase I hypersensitive element, *Hss1*, ∼5kb upstream of the *Ly49a* gene that is conserved in other *Ly49* genes and required for expression of the *Ly49a* transgene (*28*).

Our central hypothesis is that enhancers, rather than simply being permissive for RME, both limit and directly control the probability of expression of *Ly49* genes—and RME alleles generally—in a stochastic and binary fashion. Binary enhancer action, when limiting, may represent a causal mechanism of RME, explaining the pervasiveness of RME across genes and cell types. We have carried out genetic dissection and population analyses to demonstrate that enhancers control the probability of allelic expression and have provided a more general model of the role of enhancers in RME as well as in other developmentally regulated genes.

## Results

### Elements upstream of the Ly49 and Nkg2 family genes are transcriptional enhancers with activity in mature NK cells

*Ly49* family genes are expressed in a mitotically stable RME fashion by NK cells. Each harbors an accessible chromatin site (*Hss1*) ∼5 kb upstream of the transcription start site (TSS) (Fig. 1, A and B; fig. S1A). We noticed that related NK receptor genes, including the variegated *Nkg2a* gene and the *Nkg2d* gene (expressed by ∼all NK cells), harbor similar elements which we named *Nkg2a_5’E_* and *Nkg2d_5’E_* respectively (Fig. 1, A and B; fig. S1A). All *Hss1* and *5’E* elements are bound by a similar suite of factors including Runx3, T-bet, and the enhancer-associated acetyltransferase p300 (fig. S1A).

**Figure 1.**
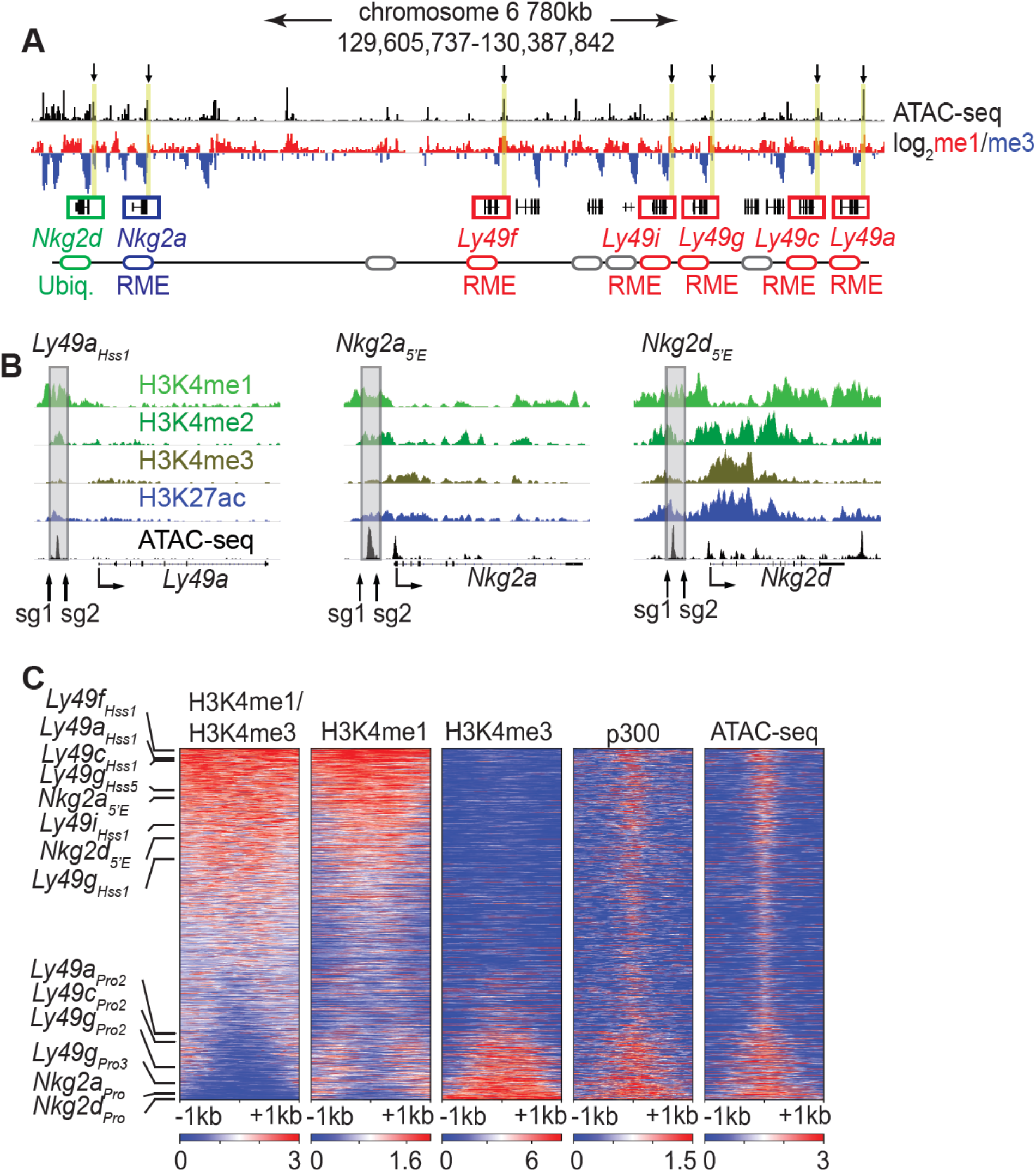
The *Ly49_Hss1_* and *Nkg2_5’E_* elements display chromatin features of enhancers. (**A**) ATAC-seq and H3K4me1:me3 log_2_ ratio ChIP-seq data of relevant NKC genes in primary NK cells; red denotes positive me1:me3 ratios (enhancer-like) while blue indicates negative values (promoter-like). Approximate gene locations are indicated (bottom). Grey ovals represent additional undiscussed *Ly49* genes. Vertical yellow bars and arrows denote the positions of the *Hss1* and *5’E* enhancers at the indicated genes. Data are sourced from ref (*55*) (**B**) Normalized ChIP-seq and ATAC-seq results (sourced from (*55*)), showing enhancer and promoter histone modifications at *Ly49a*, *Nkg2a* and *Nkg2d.* Approximate locations of sgRNAs used in this study to delete enhancers are shown. All datasets are presented with the same vertical scale across sub-panels. (**C**) Heatmaps depict 51,650 ATAC-seq peaks in primary NK cells (excluding peaks ranking in the bottom 5% for either H3K4me1 or H3K4me3) ranked according to H3K4me1:me3 ratio of average ChIP-seq signal calculated over a 2kb window centered on the ATAC-seq peak midpoint. The indicated data are displayed over these peaks in each heatmap. The locations of selected NK receptor gene *Hss1*, *5’E* and promoter elements within the me1:me3 ranking are shown. H3K4 methylation data are sourced from ref (*55*) while p300 is sourced from ref (*56*).

The *Ly49_Hss1_* elements were hypothesized to serve as upstream bidirectional promoters active only in immature, developing NK cells that switch the associated genes on or off depending on the direction of transcription (*29*). Recent evidence suggested instead that the *Ly49_Hss1_* elements are transcriptional enhancers (*30*), but this conclusion has in turn been contested (*31*). To resolve this issue, we analyzed published ChIP-seq data generated in mature primary splenic NK cells, using the H3K4me1:me3 ratio as an indicator of regulatory element identity (*32*). The *Hss1* and *5’E* elements are all enriched in H3K4me1 relative to H3K4me3 (Fig. 1A), indicating enhancer identity. The putative NK receptor gene enhancers all ranked in the top 32% of ATAC-seq accessible peaks with respect to the H3K4me1:me3 ratio. In contrast, known promoters of the respective genes ranked in the bottom 21% (Fig. 1C). In a deeper analysis, we independently defined enhancers and promoters in mature NK cells. NK cell promoters were defined as previously annotated mouse promoters from the EDPNew database (*33*) enriched in H3K27ac in NK cells, and enhancers were defined as ATAC-seq peaks bound by the p300 histone acetyltransferase that do not overlap with the promoter list. Enhancers defined in this manner were highly skewed to high H3K4me1:me3 ratios, and promoters to low ratios (fig. S1B). All *Ly49_Hss1_* and *Nkg2_5’E_* elements were classified as enhancers based on the p300-bound enhancer dataset (fig. S1B). These findings provide definitive support for the conclusion that *Ly49_Hss1_* and *Nkg2_5’E_* elements represent enhancers.

To test whether the *Ly49g_Hss1_* and *Nkg2a_5’E_* enhancers are required in mature NK cells, we adapted a CRISPR/Cas9 nucleofection protocol developed to edit primary human T cells (*34*) (fig. S2). We used NK cells from (B6 x BALB)F_1_ hybrid mice and sorted NKG2A^B6^+ or Ly49G2^B6^+ cells using B6-allele reactive monoclonal antibodies against each receptor (*25, 27*) in order to follow the fate of a single allele in each case (fig. S2A). Editing efficiencies of NK cells were lower than that of T cells, resulting in only 30% or fewer cells with disruption of the control *Ptprc* locus encoding CD45 (fig. S2B). Targeting *Nkg2a_5’E_* increased the percentage of NKG2A^B6^-negative cells from ∼10% to ∼20-40%. (fig. S2, C-E), in line with our theoretical maximum editing efficiency. Similarly, targeting *Ly49g_Hss1_* resulted in marked loss of Ly49G2^B6^ expression, with minimal (<5%) loss of expression in non-targeting or non-nucleofected (no zap) control conditions (fig. S2, F-H). These data show that the NK receptor gene enhancers play critical roles in the maintenance of active alleles in mature NK cells, and argue against the proposal that *Hss1* elements are only required in immature NK cells as once proposed (*29, 35*).

### The Ly49g_Hss1_ and Nkg2a_5’E_ enhancers are constitutively accessible

Analysis of bulk NK cells did not reveal a correlation between the gene expression frequency of an NK receptor gene and the accessibility, TF occupancy, or H3K27ac modifications of *Hss1* and *5’E* enhancers (fig. S1A). This lack of concordance raised the possibility that these enhancers were similarly active and occupied by TFs upstream of both silent and active alleles, as has been observed for RME genes genome-wide in F_1_ clones (*4, 10*). It has not previously been possible, however, to address this issue for an RME gene in freshly isolated *ex vivo* cell populations.

To purify populations of cells expressing different alleles of *Nkg2a*, we stained (B6 x BALB/c)F_1_ NK cells with allele-specific antibodies (*27*), allowing us to sort and perform ATAC-seq on NK cell populations expressing all four configurations of alleles: expressing both alleles of *Nkg2a*, only B6, only BALB, or neither (Fig. 2, A and B). SNP-split (*36*) analysis of reads demonstrated that the enhancer element *Nkg2a_5’E_* was accessible on both active and inactive alleles in all four populations, whereas the *Nkg2a* promoter was accessible only at active alleles (Fig. 2B). We used a similar allele-specific staining protocol (*25*) to sort and analyze cells expressing either, both or neither Ly49G2 allele (Fig. 2, C-D). The *Ly49g_Hss1_* enhancer was accessible on both active and inactive alleles in all four populations, whereas the dominant promoter *Pro3* (*37*) was accessible only on the active allele (Fig. 2D). Notably, the *Ly49g* gene harbors a second minor enhancer element, *Ly49g_Hss5_* (Fig. 1C; fig. S1B), which was similarly accessible at all alleles (Fig. 2D).

**Figure 2.**
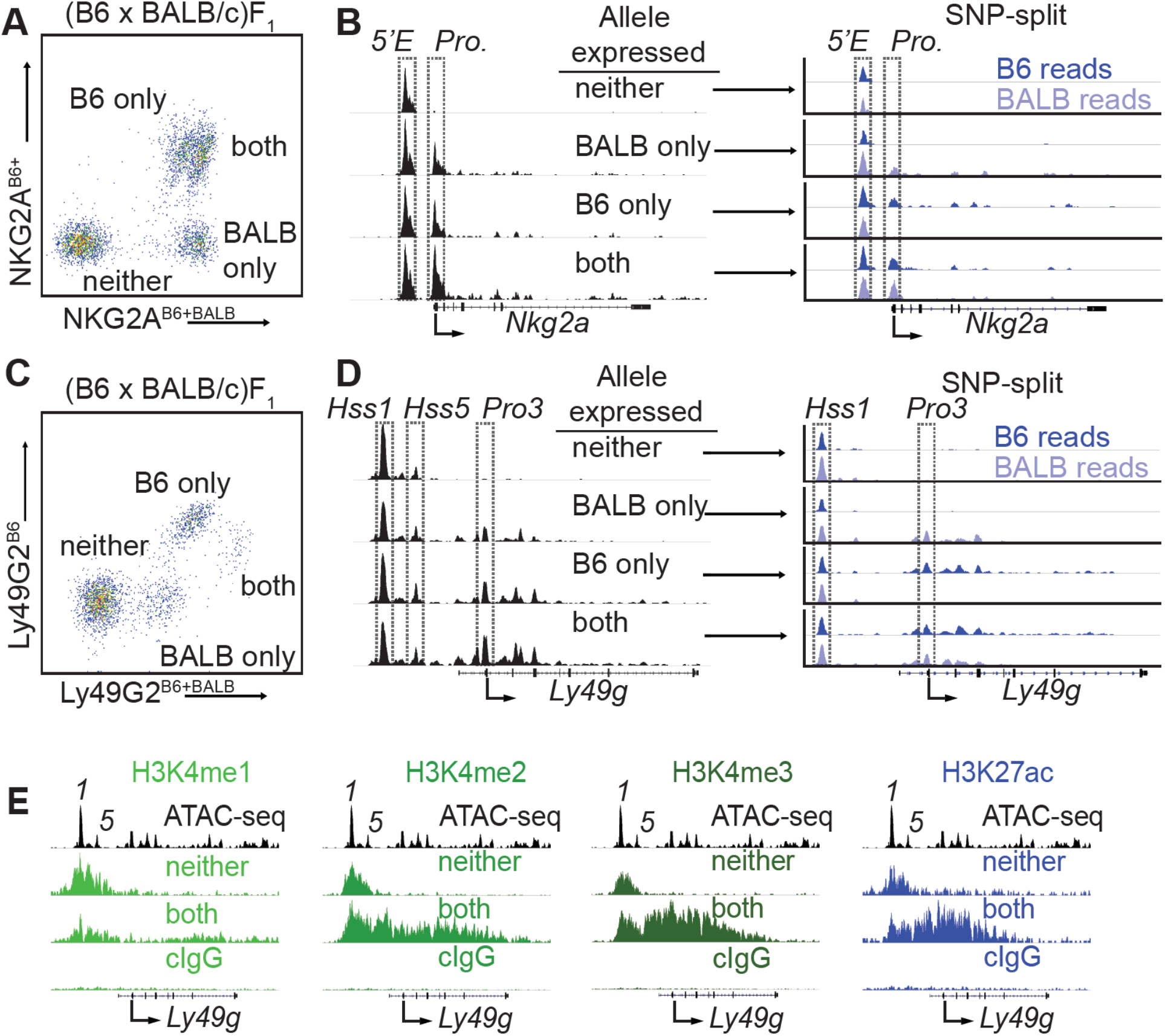
*Nkg2a_5’E_* and *Ly49g_Hss1_* are constitutively accessible, while promoters are accessible only at expressed alleles. (**A**) FACS plot depicting splenic NK cells from a (B6 x BALB/c)F_1_ hybrid mouse stained with allele-specific antibodies, allowing separation of NK cells expressing both, either, or neither NKG2A allele. (**B**) (left) Normalized ATAC-seq data generated from the 4 cell populations depicted in (A) aligned to the mm10 reference genome. (right) Allele-informative reads were binned according to chromosome of origin, and displayed as signal mapping to the B6 or BALB/c chromosome. The *Nkg2a_5’E_* enhancer and promoter (*Pro.*) are boxed (dotted line) (**C** and **D**) Data are as in (A and B), but using an allele-specific staining protocol with respect to the Ly49G2 receptor. *Ly49g_Hss1_*, *Ly49g_Hss5_* and the dominant TSS (*Pro3*) are boxed. (**E**) CUT&RUN data depicting each of 4 indicated histone modifications, of the *Ly49g* gene in IL-2 expanded NK cells sorted to express neither “N” or both “B” alleles of *Ly49g*, or a control mouse IgG2a*κ* (cIgG) antibody in IL-2 expanded NK cells. The ATAC-seq pattern is shown for reference above each analysis; *Hss1* is denoted as *“1*”, Hss5 is denoted as “*5*”. Arrows depict the locations of the dominant *Pro3* TSS. All ATAC-seq and CUT&RUN data within a sub-panel are presented with the same vertical scale.

These data demonstrated that enhancers within the *Ly49* and *Nkg2* gene families behave similarly to those of other RME genes analyzed in F_1_ hybrid clones (*4, 10*), exhibiting an accessible configuration whether or not the gene was expressed. Importantly, this analysis further validated the NK receptor genes as a model for RME. While initially surprising, the decoupling of enhancer and promoter accessibility seen at NK receptor genes and other RME loci is consistent with a binary model of enhancer action, where enhancer activation occurs in all cells of a given type and acts stochastically to raise the binary “on or off” probability of gene expression, rather than regulate the per-cell amount of expression (*11, 18*).

We extended these observations by analyzing the pattern of active enhancer associated marks at silent and active alleles of *Ly49g.* We sorted IL-2-expanded NK cells expressing neither (N) or both (B) Ly49G2 alleles from (B6 x BALB)F_1_ mice and performed CUT&RUN for the enhancer-associated H3K4me1/2/3 and H3K27ac modifications. The *Ly49g* promoter and gene body displayed striking enrichment of H3K4me2/3 and H3K27ac in cells that expressed both *Ly49g* alleles, and as predicted lacked these modifications in cells where neither allele was expressed (Fig. 2E). Notably, the *Ly49g_Hss1_* and *Ly49g_Hss5_* enhancers displayed equal enrichment of H3K27ac in cells expressing both alleles or neither (Fig. 2E). As H3K27ac delineates active as opposed to poised enhancers (*32*), these data suggest constitutive enhancer activation on both silent and active alleles.

### Ly49a_Hss1_ and Nkg2a_5’E_ are required for gene expression in vivo, and act in cis

We tested the requirement for *Ly49a_Hss1_* in the endogenous locus by deleting it in the B6 germline via CRISPR/Cas9 editing (Fig. 3, A-C, fig. S3, A and B). *Ly49a^Hss1^*^Δ^*^/Hss1^*^Δ^ mice completely lacked Ly49A expression (Fig. 3, B and C; fig. S3B), but importantly expression of other Ly49 receptors was unaffected (fig. S3B), supporting the notion that the variegated NK receptor genes are regulated proximally and independently of each other. *Ly49a*^+*/Hss1*Δ^ mice displayed an intermediate percentage of Ly49A+ cells (Fig. 3, B and C; fig. S3B).

**Figure 3.**
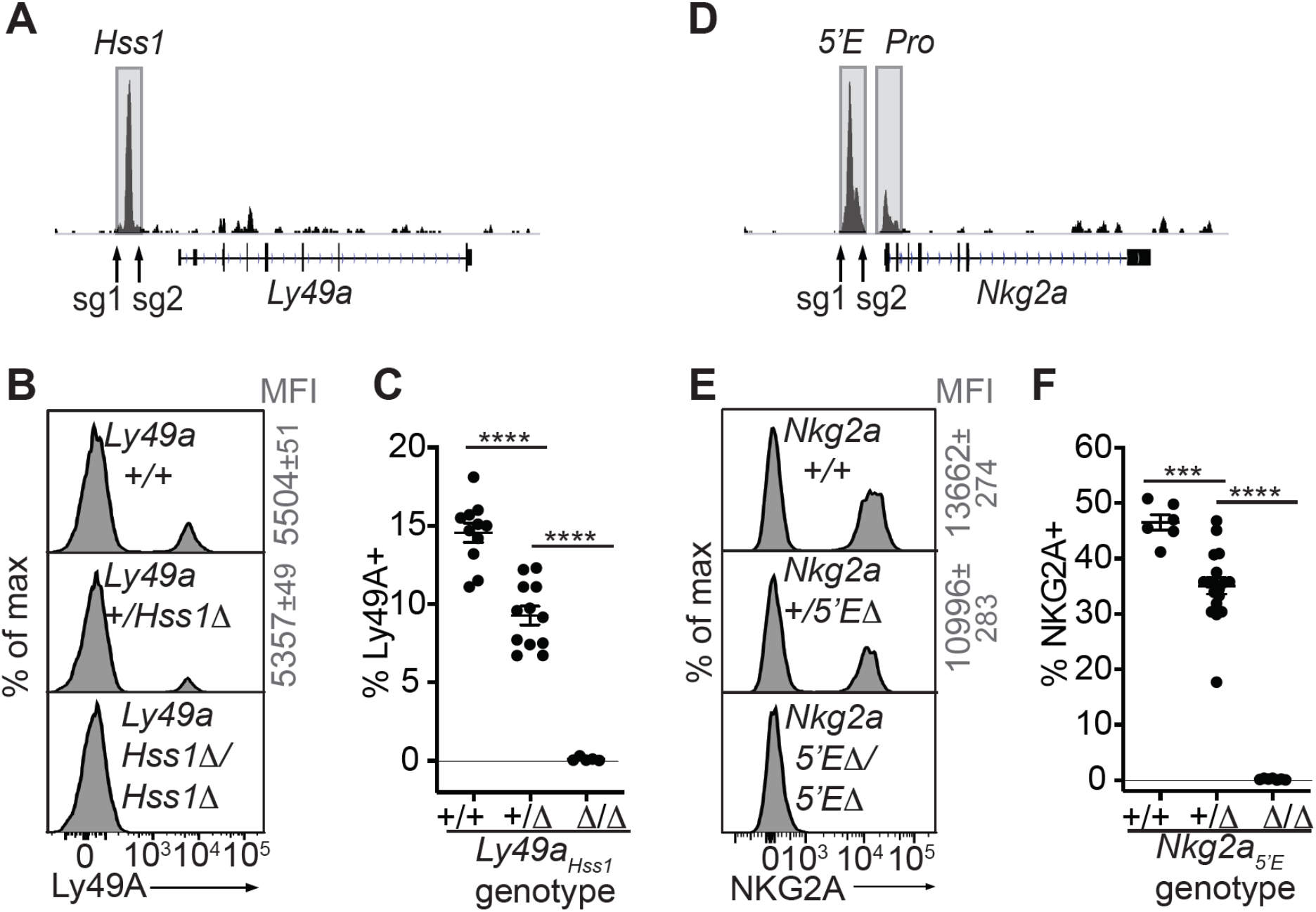
The *Ly49a_Hss1_* and *Nkg2a_5’E_* enhancers are required for gene expression. (**A**) Locations of sgRNAs used to delete *Ly49a_Hss1_* in the B6 germline; NK cell ATAC-seq are displayed for reference. (**B-C**) Ly49A staining of the indicated *Ly49a_Hss1_* deletion littermates. MFI of staining + SEM are depicted in grey. In (C), data are combined from two independent experiments (means±SEMs, *n*=5-12). (**D-F**) Data as in (B-C) for *Nkg2a_5’E_*. Data in (F) are combined from two experiments with the *Nkg2a_5’E_(B3Δ)* allele (fig. S3C) and were recapitulated in analysis of the *B1Δ* allele (means±SEMs, *n*=6-18). \*\*\*\**P* <0.0001; \*\*\**P* <0.0001 using One-way ANOVAs with Tukey’s multiple comparisons.

As with *Ly49a_Hss1_*, deletion of both allelic copies of *Nkg2a_5’E_* in the germline eliminated NKG2A expression, and heterozygous mice displayed a reduced frequency of NKG2A+ NK cells (Fig. 3, D and E; fig. S3, C and D).

Whether the activity of constitutively accessible enhancers of RME genes is coordinated in *trans* via a dedicated epigenetic mechanism that enacts RME is not known. We addressed the *cis* vs *trans* activity of *Nkg2a_5’E_* and *Ly49a_Hss1_* in F_1_ hybrids using allele-discriminating antibodies. F_1_ hybrid mice between B6-*Nkg2a^B6-5’E^*^Δ^ mice and BALB/c mice were generated (*Nkg2a^B6-5’E^*^Δ^*^/BALB/c+^* heterozygotes) along with WT F_1_ littermates. The percentages of cells expressing all four combinations of NKG2A alleles were determined using allele-specific NKG2A antibodies we previously generated. Assuming that the *Nkg2a_5’E_* acts only in *cis* we calculated the expected changes in the frequencies of these cells in the heterozygotes. The experimental data closely mirrored the predictions (fig. S4, A-C). Therefore, the constitutively accessible *Nkg2a_5’E_* acted *in cis* and independently of the activity of the other copy. Similarly, in *Ly49a^B6-Hss1^*^Δ^*^/Balb/c+^* heterozygotes, the BALB/c allele was unaffected when expression of the B6 allele was abrogated (fig. S4, D-F).

### A cis-acting secondary enhancer in the Ly49g locus contributes to the high expression frequency of Ly49G2

Both the *Ly49a* and *Nkg2a* gene loci harbor only a single prominent proximal enhancer-like site (Fig. 1, A and B), and are completely dependent on those enhancers for expression (Fig. 3), complicating analysis of the role of *Ly49a_Hss1_* and *Nkg2a_5’E_* in the RME of their target genes. We reasoned that analysis of an RME NK receptor gene with multiple proximal enhancers could reveal the role of overall enhancer strength in regulating expression frequency. We hypothesized, in accordance with the binary model, that despite the presence of multiple (weak) enhancers, enhancer activity at such loci is limiting resulting in RME. We predicted that limiting enhancer activity further by deleting a secondary enhancer in a natural RME gene would reduce, but not abrogate, gene expression probability.

The *Ly49g* locus is expressed by ∼50% of NK cells and contains both an *Hss1* element and another constitutively accessible enhancer, *Ly49g_Hss5_* (Fig. 2D). Interestingly, the corresponding region of the highly related *Ly49a* gene, which is expressed by only ∼17% of NK cells, is much less accessible and presumably less active (Fig. 4A).

**Figure 4.**
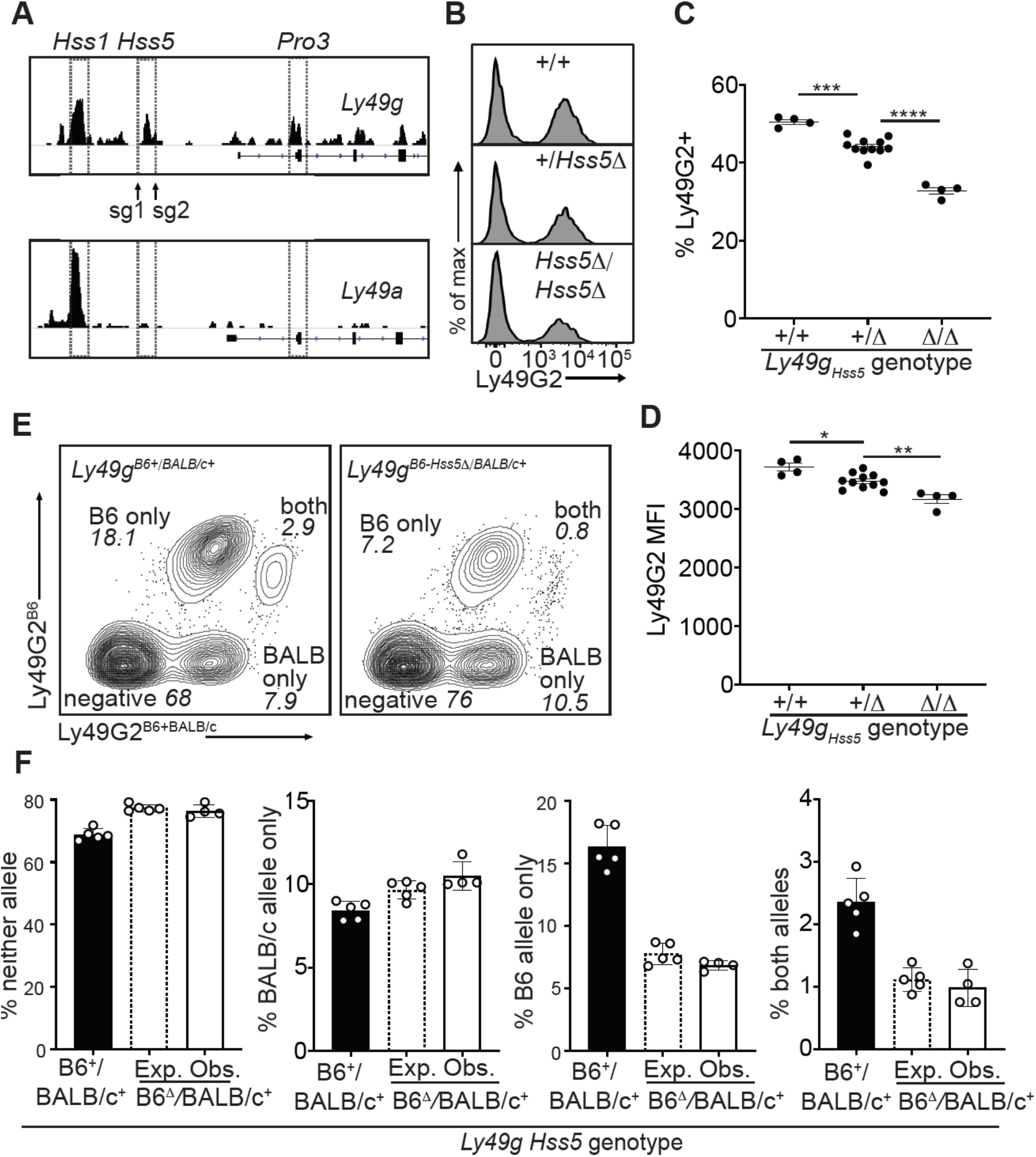
A minor cis-acting enhancer amplifies Ly49G2 expression frequency. (**A**) Normalized ATAC-seq tracks of *Ly49a* and *Ly49g* in bulk NK cells; data are on the same vertical scale (top and bottom). *Hss1* and *Hss5* enhancers and the *Pro3* promoter are highlighted. sgRNAs used to generate *Ly49g_Hss5_*_Δ_ alleles are shown (arrows). (**B**) Ly49G2 staining of NK cells in the indicated *Ly49g _Hss5_* deletion littermates (B2Δ allele, fig. S5A). (**C-D**) Ly49G2 percentages (C), and mean fluorescence intensities of the positive populations (D) (*n*=4-11). Similar results were obtained with the B1Δallele (fig. S5B). (\**P* <0.05; \*\**P* <0.01; \*\*\**P* <0.001; \*\*\*\**P* <0.0001 using One-way ANOVA with Tukey’s multiple comparisons). (**E**) Flow cytometry plots of gated *Ly49g^B6-Hss5^*^Δ/*BALB/c+*^ NK cells using Ly49G2^B6^ specific and Ly49G2^B6+BALB/c^ specific antibodies (right) and a wildtype littermate (left). (**F**) Expected and observed percentages of populations depicted in “E” in F1 mice with the *Hss5Δ* (hatched bar is expected, white bar is observed) or wildtype (black) *Ly49g* allele. Expected frequencies were calculated assuming stochastic *cis* regulation of alleles (see Methods; note effect of genetic background). Data are representative of two experiments. All error bars represent SEM.

Germline deletion of *Ly49g_Hss5_*, in homozygous configuration, resulted in a depressed percentage of Ly49G2+ cells (35%) compared to WT mice (50%), with only a minor change in expression per cell (measured by mean fluorescence intensity of staining, MFI) (Fig. 4, B-D; fig. S5, A and B). Heterozygous mice displayed an intermediate percentage of Ly49G2+ cells (Fig. 4, B and C; fig S5B). To test whether *Ly49g_Hss5_* acts entirely *in cis*, we crossed *Ly49g^Hss5^*^Δ^ to BALB/c mice. The NK cell populations in the F_1_ mice expressing the Ly49G2^B6^ alleles were reduced, and the populations expressing neither allele or only Ly49G2^BALB/c^ were increased, in the proportion expected under probabilistic action of *Hss5 in cis* (Fig. 4, E and F). Thus, the constitutively active enhancer *Ly49g_Hss5_* is directly involved in regulating Ly49G2 expression frequency, and explains, at least in part, the high expression frequency of Ly49G2 in relation to other receptors including Ly49A.

### Deletion of Nkg2d_5’E_ is sufficient to recapitulate stable RME in Nkg2d

Our hypothesis that RME of NK receptor genes is imparted by limiting binary enhancer activity predicts that a receptor expressed by all NK cells may be converted into a variegated receptor by weakening enhancer activity, for example by deleting one of multiple associated enhancer elements. We tested this for the *Nkg2d* gene encoding the NKG2D immunostimulatory receptor, which is expressed by all NK cells (*38*), is distantly related to the *Nkg2a* and *Ly49* genes, and is flanked on both sides by enhancer-like chromatin, suggesting possible regulation by multiple enhancers (Fig. 1B). Deletion of the enhancer-like ATAC-accessible site ∼5 kb upstream of the *Nkg2d* gene (*Nkg2d_5’E_*) (Fig. 1B; fig. S5, C and D), resulted in variegated NKG2D expression in *Nkg2d^5’E^*^Δ^*^/5’E^*^Δ^ animals (Fig. 5, A and B). Only ∼65% of NK cells expressed NKG2D in *Nkg2d^5’E^*^Δ^*^/5’E^*^Δ^ animals. The expression level per cell was only modestly affected and to an extent consistent with largely monoallelic expression vs expression in some cells of both alleles (Fig. 5C and fig. S5E). These data suggest that the primary role of *Nkg2d_5’E_* is to increase the probability rather than the degree of *Nkg2d* expression, in line with the binary model of enhancer action.

**Figure 5.**
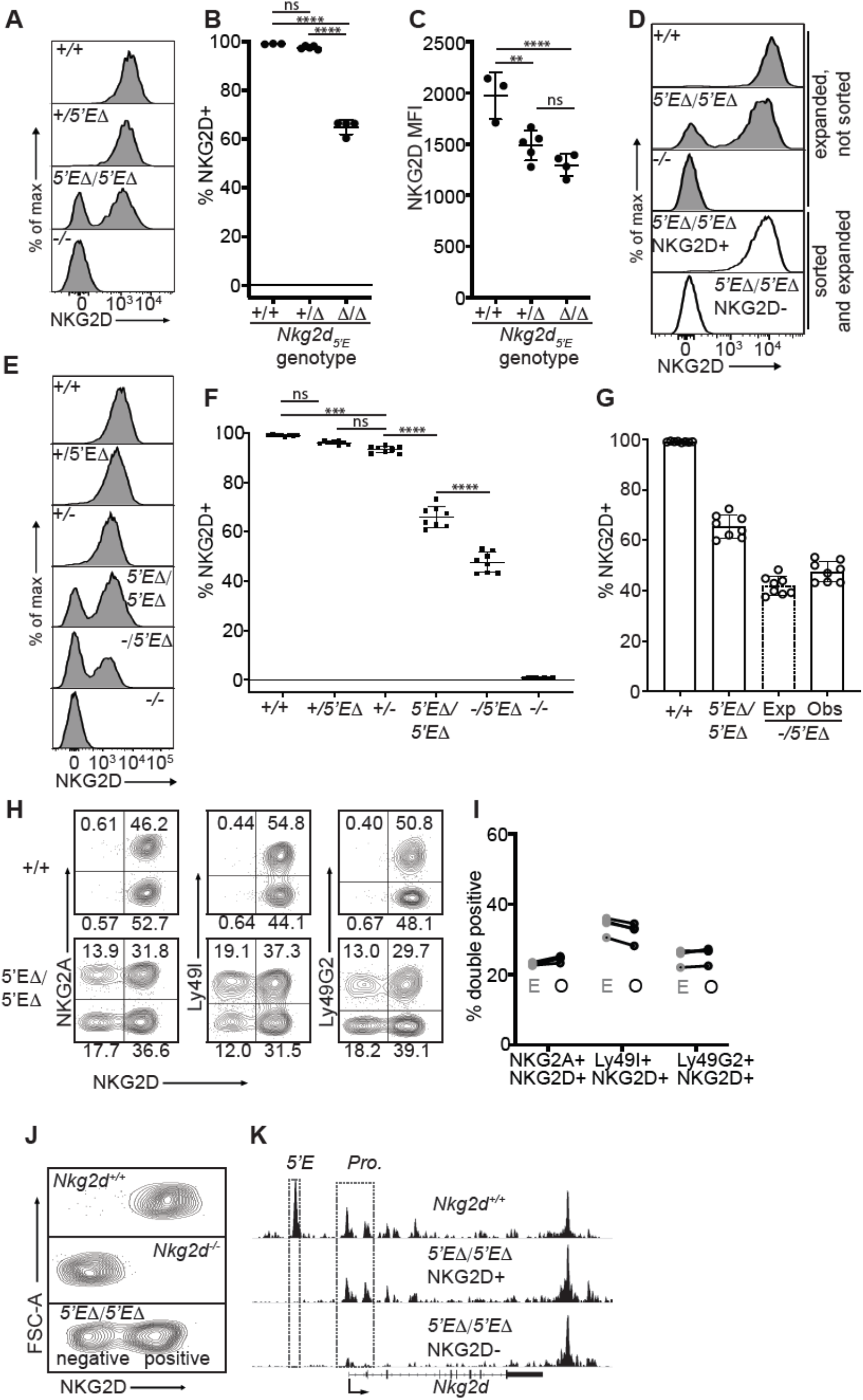
*Nkg2d_5’E_* deletion results in mitotically stable RME, fully recapitulating natural variegation. (**A-C**) NKG2D staining of splenocytes from *Nkg2d_5’E_* deletion littermates (B1Δ allele), and an *Nkg2d^-/-^* mouse (*n*=3-5). Results are representative of four experiments with two deletion alleles (fig. S5, C and D) (**D**) Splenocytes from *Nkg2d^5’E^*^Δ^*^/5’E^*^Δ^ mice were cultured with IL-2 for 2-3 days before sorting NKG2D+ and NKG2D-NK cells, which were expanded in fresh IL-2 medium for 8-10 days before analysis (white fill). Expanded, unsorted NK cells in grey. (**E**) Staining of splenic NK cells from mice of six genotypes. “+”, “-” and “Δ” refer to wildtype, gene knockout, and *5’E* deletion alleles, respectively. (**F**) Quantified results in (E) compiled from two experiments. (**G**) Expected and observed percentages of NKG2D+ NK cells in *Nkg2d^5’E^*^Δ^*^/-^* mice. Expected expression is calculated based on observed NKG2D^+^ percentages in *Nkg2d^5’E^*^Δ^*^/5’E^*^Δ^ mice, assuming stochastic expression (see Methods). (**H**) Stochastic co-expression of NKG2D and NKG2A, Ly49I or Ly49G by NKp46+ NK cells in *Nkg2d^5’E^*^Δ^*^/5’E^*^Δ^ mice. WT (+/+) mice are shown for comparison. (**I**) Expected (“E”) and observed (“O”) percentages of cells coexpressing the indicated receptors in *Nkg2d^5’E^*^Δ^*^/5’E^*^Δ^ mice. Expected percentages were calculated by mutiplying percentages of cells in each mouse expressing each receptor individually (*n*=4). Data are representative of two experiments. (**J**) NKG2D staining of presorted gated NK cells from *Nkg2d^5’E^*^Δ^*^/5’E^*^Δ^ mice (bottom), compared to wildtype and *Nkg2d^-/-^* NK cells. (**K**) Normalized ATAC-seq tracks generated from NKG2D+ and NKG2D-cells sorted from the *Nkg2d^5’E^*^Δ^*^/5’E^*^Δ^ mouse shown in (J) and are presented on the same vertical scale. ATAC-seq results for WT splenic NK cells were sourced from (*55*) and auto-scaled to match the data generated from the *Nkg2d^5’E^*^Δ^*^/5’E^*^Δ^ mouse. Error bars represent SEM. \*\**P* <0.01; \*\*\**P* <0.001; \*\*\*\**P* <0.0001, computed using One-way ANOVAs with Tukey’s multiple comparisons.

Significantly, the expression state of *Nkg2d^5’E^*^Δ^ alleles was mitotically stable. NK cells from the enhancer knockouts were stimulated with IL-2 for 2-3 days before sorting NKG2D+ and NKG2D-populations, and further expanded in IL-2 for an additional 8-10 days, where they underwent an ∼10-100 fold expansion. The NKG2D+ and NKG2D-phenotypes were highly stable despite extensive proliferation (Fig. 5D).

In heterozygotes with the *Nkg2d^5’E^*^Δ^ allele on one chromosome and an *Nkg2d* knockout allele on the other (*-/5’EΔ*) the percentage of NKG2D+ cells was lower than in *5’EΔ*/*5’EΔ* mice (Fig. 5, E and F). This nearly matched the expected percentage under the assumption that the *5’EΔ* alleles are independently regulated in the heterozygotes, i.e., the positive cells include cells expressing both alleles with a frequency that is the product of the individual frequencies (*22*) (Fig. 5G). NKG2D expression per NKG2D+ cell in *Nkg2d^5’E^*^Δ/*5’E*Δ^ animals appeared slightly higher than in *Nkg2d^+/-^* animals, consistent with a proportion of cells expressing both *Nkg2d* alleles, a feature characteristic of natural RME (*1, 6*) (Fig. 5E and fig. S5E). Together, these data strongly argue that expression of *Nkg2d^5’E^*^Δ^ alleles follows a stochastic RME pattern.

### The Nkg2d_5’EΔ_ allele mimics the expression and accessibility features of naturally variegated NK receptor genes

The stable RME of *Nkg2d^5’E^*^Δ^ alleles recapitulated the stochastic expression pattern of naturally variegated NK receptor genes. Expression of NKG2D in *Nkg2d^5’E^*^Δ/*5’E*Δ^ mice was approximately randomly distributed with respect to the naturally variegated NKG2A, Ly49G2 or Ly49I (Fig. 5H). Indeed, the co-expression frequencies approximated the products of the separate frequencies of the receptors studied (the “product rule” (*22*)) (Fig. 5I).

To examine the chromatin accessibility of the *Nkg2d* locus in *Nkg2d^5’E^*^Δ/*5’E*Δ^ mice, we performed ATAC-seq with NKG2D+ and NKG2D-cells sorted from *Nkg2d^5’E^*^Δ/*5’E*Δ^ mice (Fig. 5J). Robust promoter accessibility was detected in NKG2D+ cells but not in NKG2D-cells (Fig. 5K). The 5’E element was deleted in both populations and therefore not accessible, but a 3’ enhancer-like element was equally accessible in both populations. This accessibility pattern mirrors that of the naturally variegated NK receptor genes (Fig. 2) and RME broadly (*4*).

The results of our experiments with the *Nkg2d* gene established that stable RME and the stochastic and variegated NK receptor expression pattern could be recapitulated in full by weakening enhancer activity at a gene normally expressed in ∼all NK cells. Furthermore, the similarity of *Nkg2d^5’E^*^Δ/*5’E*Δ^ and natural NK receptor gene variegation suggests that previous examples of enhancer deletion-associated variegation such as that seen in the *Cd8a* locus (*16, 17*) are rooted in similar mechanisms as naturally-occuring RME.

### Silent NK receptor gene alleles lack repressive histone modifications associated with polycomb and heterochromatic repression

We investigated histone modifications associated with gene repression to search for clues regarding the maintenance of the active and silent epigenetic states. We assayed the polycomb-associated marks H3K27me3 and H2AK119Ub1 (H2AUb1) and the heterochromatin-associated H3K9me3, which have previously been found at inactive alleles of some other monoallelically expressed genes, notably the odorant receptors and protocadherins (*1, 4, 6, 8, 9*). CUT&RUN analysis of repressive modifications in IL-2 expanded primary NK cells that were sorted as Ly49G2-negative (expressing neither allele, designated “N”), revealed that all three modifications were prevalent in the *Hoxa* gene cluster, as expected, but the entire NKC lacked appreciable signal for any of the modifications (Fig. 6A). None of the 3 marks were enriched above background on silent *Ly49g* alleles (Fig. 6B). Importantly, many other genes associated with non-NK cell lineages (e.g., *Cd19* and *Mstn*, expressed in B cells and myocytes, respectively) were also not enriched for these repressive modifications (Fig. 6B). In contrast, other genes such as *Pdcd1* (encodes PD-1) and *Spi1* (encodes the macrophage and B-cell lineage-regulating transcription factor PU.1) displayed all 3 marks. Therefore, with respect to repressive chromatin marks, silent *Ly49g* alleles resembled several genes normally expressed in other hematopoietic cell lineages but not in NK cells, rather than known repressed genes.

**Figure 6.**
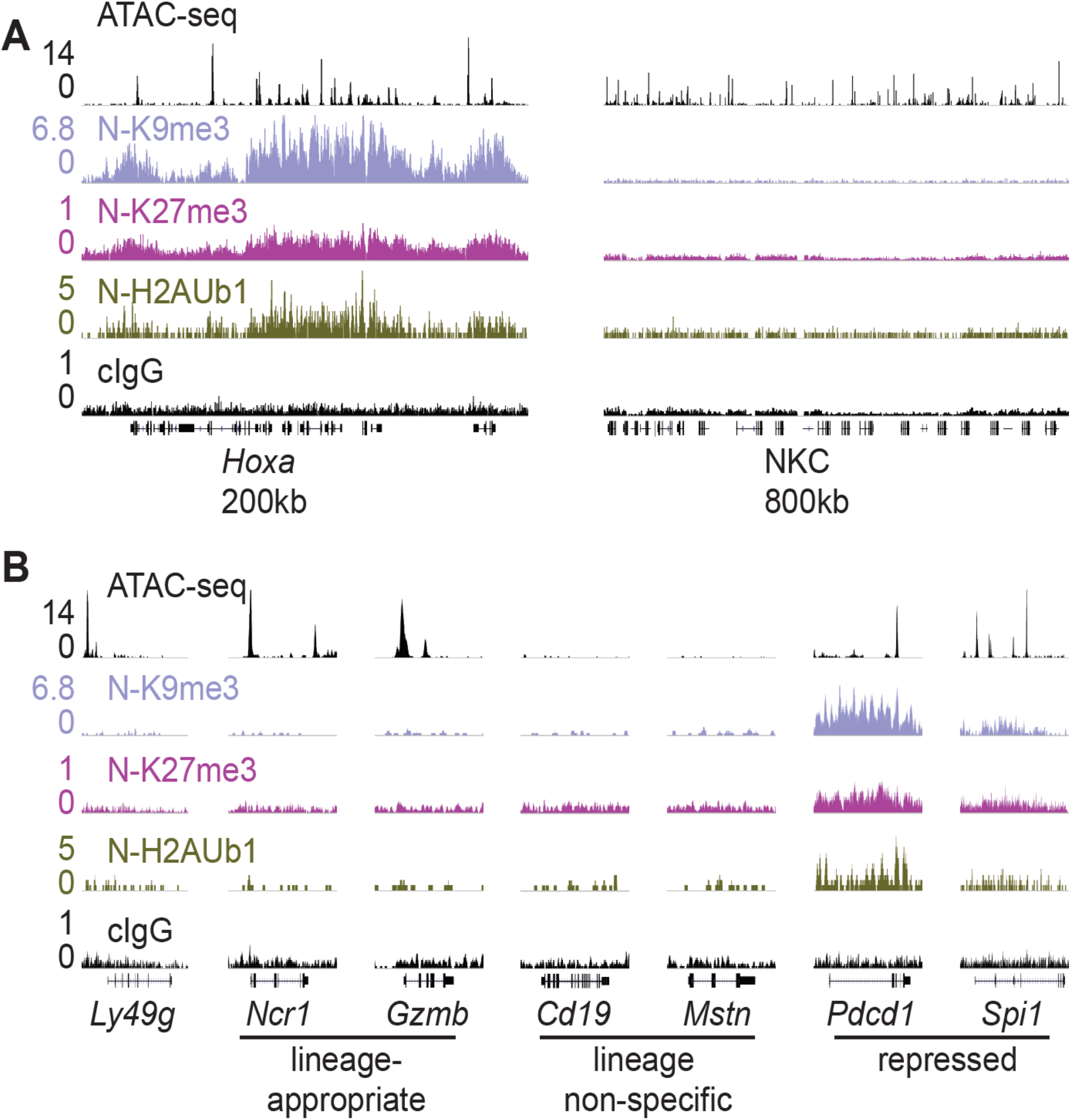
Silent NK receptor gene alleles resemble inactive genes expressed in non-NK lineages, rather than repressed genes. (**A**) Repressive histone modification CUT&RUN data generated in primary IL-2 expanded NK cells sorted to express neither allele of Ly49G2 (“N” cells). IGV screenshots depicting the indicated histone modification or analyses with control mouse IgG2a*κ* (cIgG), which binds protein A. The *Hoxa* gene cluster (left) serves as a positive control. The entire NKC gene cluster is displayed on the right. The vertical scales, indicated on the left of the panels, were matched for each type of mark for all samples analyzed and were chosen to provide strong signals for the positive control Hoxa cluster. The cIgG data were scaled the same as the H3K27me3 data, which had the weakest signal of the marks analyzed. (**B**) Data are displayed as in (A), at *Ly49g* (left), and gene loci belonging to the following classes: NK cell lineage-appropriate, NK cell lineage non-specific, and loci repressed in NK cells.

As silent *Ly49g* alleles appeared more similar to lineage non-specific genes in our analysis of repressive chromatin marks, we extended the analysis of chromatin states using ChromHMM, which integrates multiple datasets to classify the genome into subdomains based on their chromatin signatures (*39*). Using data from cells expressing neither or both Ly49G2 alleles, we constructed a minimal 3 state model corresponding to transcriptionally active chromatin (high levels of H3K27ac and H3K4me3, both active marks), repressed chromatin (H2AUb1 and H3K9me3, both repressive marks), and inactive chromatin (lacking these active or repressive marks) (fig. S6, A and B). As expected, the promoters of lineage-appropriate genes expressed in NK cells (e.g., *Ncr1, Nk1.1, Ifng*) fell into the “active” chromatin state 1 (fig. S6C). Notably, genes commonly regarded as markers of non-NK cell hematopoietic lineages (e.g., *Cd3e, Cd19, Ly6g, Siglech*) fell into the “inactive” chromatin state 2 (fig. S6D). Finally, promoters of other genes, often encoding transcription factors that promote non-NK cell fates such as *Bcl11b*, *Batf3* and *Pax5,* fell into the “repressed” state 3 (fig. S6E). These data suggest that many genes encoding immune effector molecules associated with non-NK lineages are not actively repressed but are inactive and stably silent, whereas genes promoting non-NK cell fates are actively repressed.

In cells expressing both copies of *Ly49g*, the enhancer, promoter and gene body all fell within the active state 1 (fig. S6F), whereas in cells expressing neither copy, the enhancer remained in the active state 1 but the promoter and gene body became inactive (state 2) rather than repressed (state 3). Indeed, it was striking that the NKC as a whole lacked repressive state 3 chromatin (Fig. 6A; fig. S6F). The lack of the repressive chromatin state at silent NK receptor genes suggests that repressive chromatin may not be required for stable RME generally, potentially explaining why repressive chromatin signatures are not a consistent feature of silent RME alleles in other instances (*1, 6, 8, 9*). In lieu of active repression, other mechanisms must be invoked for the maintenance of RME patterns through cell division.

### Mitotically stable RME is likely far more common than previously appreciated

Our findings that RME is rooted in broad and probabilistic properties of gene activation raised the possibility that these principles might apply to many and perhaps all genes. We hypothesized that many genes exhibit a minor extent of RME but escaped previous detection due to the methods used, which were limited by clone numbers and therefore lacked the resolution to detect very rare monoallelic expression (*3, 5, 8, 9*). We employed flow cytometry to analyze millions of primary cells *ex vivo* for rare monoallelic expression of several membrane proteins, starting with NKG2D. We noticed that ∼2.5% of NK cells in *Nkg2d^+^*^/*-*^ mice lacked expression of NKG2D, whereas the percentage in *Nkg2d^+^*^/*+*^ mice was close to 0% (Fig. 5F; Fig. 7, A and B). These data suggested that rare monoallelic expression of WT *Nkg2d* was obscured by expression of at least one allele in nearly all NK cells. Indeed, assuming allelic independence, the failure of each allele to be expressed in a random 2.5% of all NK cells (from here on refered to as an “allelic failure rate”) translated to only 0.063% of cells lacking both alleles. Importantly, NKG2D-cells in both *Nkg2d^5’E^*^Δ/*5’E*Δ^ and *Nkg2d^+^*^/*-*^ mice were as likely to express NKG2A, Ly49G2 or Ly49I as NKG2D+ cells, suggesting stochastic expression of the WT *Nkg2d* allele (fig. S7, A-C). Thus, even the WT *Nkg2d* gene is expressed in an RME fashion.

**Figure 7.**
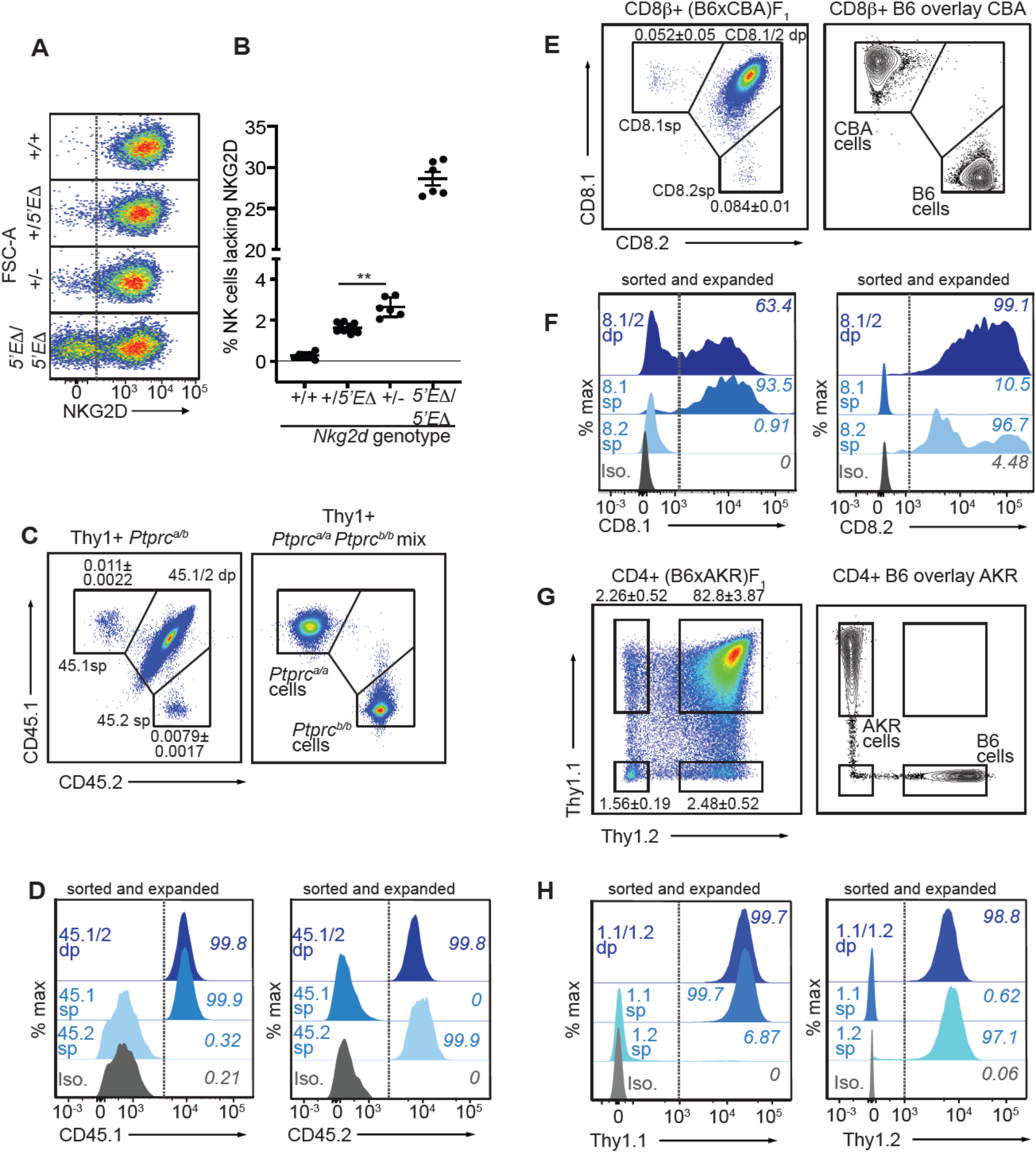
*Nkg2d, Ptprc*, *Cd8a and Thy1* are RME genes. (**A, B**) Flow cytometry (A) and quantification of %NKG2D-negative cells (B) of selected *Nkg2d* genotypes. *P*=0.002, student’s *t*-test. (**C**) Monoallelic CD45 expression. Flow cytometry of gated Thy1+ cells pooled from 2 *Ptprc^a/b^* mice (left). The mean percentages±SEM of each monoallelic population, combined from 3 experiments, are depicted within the plot. Right panel: a mixture of cells from *Ptprc^a/a^* and *Ptprc^b/b^* mice. (**D**) CD45 allele single positive and double positive T cell populations were sorted from *Ptprc^a/b^* mice using gates in panel C, expanded for 1 week *in vitro*, resorted to purity and expanded an additional ∼5-8 fold. Histograms show CD45.1 and CD45.2 staining for the sorted populations after expansion. (**E-F**) Monoallelic expression of CD8*α* in (B6 x CBA)F1 mice presented as in (C) and (D). (F) shows CD8*β*+ cells from F1 mice sorted and expanded twice as in (D) (∼5-10 fold expansion in the second stumulation). (**G-H**) Data are displayed as in (C-F) but with respect to Thy1 allelic expression on CD3+CD4+ T cells in (B6 x AKR)F_1_ hybrid mice (6-8 fold expansion in the second stimulation). All experiments are representative of 2-3 performed. Error bars represent SEM.

To extend this approach, we sought to analyze allelic expression of receptor genes for which A) allele-specific antibodies exist and, B) expression is normally considered to be universal in defined lymphohematopoietic lineages. We first analyzed expression of allelic variants of the *Ptprc* gene encoding CD45, a membrane phosphatase expressed by all lymphohematopoietic cells. The *Ptprc^a^* allele, encoding CD45.1, and the *Ptprc^b^* allele, encoding CD45.2, are easily discriminated by flow cytometry with monoclonal antibodies in congenic mice. Heterozygous *B6-Ptprc^a/b^* mice are expected to express both alleles on all B or T cells, but we were able to detect clearly defined, albeit very rare (∼0.01%) subpopulations of B or T cells expressing only one allele or the other (Fig. 7C; fig. S8, A and B). The monoallelic cells exhibited similar staining intensity as homozygous *B6-Ptprc^a/a^* and *B6-Ptprbc^b/b^* cells analyzed in parallel. Given the very low frequency of monoallelic *Ptprc* expression, cells lacking CD45 altogether would be predicted to be extremely rare and indeed were not detected. Sorted CD45.1 and CD45.2 single positive T cells from *B6-Ptprc^a/b^* mice retained monoallelic expression over 5-8 fold expansion after stimulation with CD3/CD28 beads *in vitro*, demonstrating that RME of *Ptprc* is mitotically stable (Fig. 7D). Sanger sequencing of reverse-transcribed and amplified RNA isolated from the expanded cells displayed in Fig. 7D revealed that the monoallelic populations expressed only the allele detected by cell surface staining, demonstrating that the rare observed RME of *Ptprc* reflected transcriptional differences (fig. S8C). These data argued strongly against the possibility that the monoallelic cells arose due to somatic mutations in one or the other *Ptprc* allele, since most such mutations would not be predicted to disrupt transcription.

Similar analysis of *Cd8a* for RME employed allele-specific CD8*α* antibodies. For *Cd8a*, we gated on cells expressing CD8*β*, the partner chain of CD8*α*. In (B6 x CBA)F_1_ mice, approximately 0.1% of CD8*β*+ cells lacked one or the other of the CD8*α* alleles, CD8.1 or CD8.2 (Fig. 7E; fig. S8, D and E). Sorted single positive cells that retained expression of CD8β after stimulation and expansion retained expression of the initially selected allele of CD8*α*, demonstrating mitotically stable RME (Fig. 7F). Finally, analysis of *Thy1,* which is thought to be expressed by all T cells in mice, also employed allele-specific antibodies to discriminate the allelic Thy1.1 and Thy1.2 proteins, expressed by AKR and B6 strains, respectively. In (B6 x AKR)F_1_ hybrids ∼5% of CD4+ T cells expressed only one allele or the other (Fig. 7G; fig. S8, F and G). Again, the monoallelic populations displayed an impressive degree of mitotic stability (Fig 7H). Hence, Thy1 represents an RME gene with a remarkably high allelic failure rate. In conclusion, RME was detectable for all four genes we examined, all of which were previously considered to be expressed in all cells of the lineages analyzed.

These findings supported the notion that RME is characteristic of many genes and is a natural consequence of enhancer-promoter interactions, rather than a specialized form of gene expression. We quantified allelic failure rates of the various genes examined in this study (fig. S8H). We propose that in the absence of selection for biallelic expression, most genes exist along a continuum of allelic failure rates, and that RME and “non-RME” genes differ quantitatively with respect to allelic failure rather than qualitatively with respect to dedicated, RME-specific regulatory programs.

## Discussion

The *Ly49a Hss1* element was previously reported to be a “switch” element active only in immature NK cells (*29*). Our extensive analysis herein demonstrated that *Hss1* displays properties of enhancers in mature cells, consistent with the conclusions of others based on reporter analysis (*30*). Furthermore, the loss of Ly49G2 expression after deletion of *Ly49g_Hss1_* that we have documented in mature Ly49G2+ cells is inconsistent with a solely developmental role of *Hss1*. Finally, our results showed that variegation arises and is modulated by enhancer deletion (including *Ly49g_Hss5_* and *Nkg2d_5’E_*) rather than introduction of variegating switch elements.

The main significance of our results is to link enhancer deletion-associated variegation with naturally-occuring RME and place previous results in the context of a pervasive biological phenomenon. Remarkably, deletion of an enhancer upstream of the *Nkg2d* gene imparted an RME expression pattern that fully recapitulated the stochasticity, mitotic stability and promoter accessibility features of naturally variegated receptor genes. In this instance, enhancer-like elements downstream of *Nkg2d* may suffice to impart the lower frequency of expression. The striking commonalities of enhancer deletion variegation and natural variegation of NK receptor genes and other RME genes argues that RME is an extreme manifestation of the inherent probablistic nature of stable gene activation rather than a specialized mechanism to impose a variegated expression pattern.

The data also reveal the quantitative impact of enhancer strength on allelic expression frequencies. Deletion of *Ly49g_Hss5_*, a relatively minor and constitutively accessible enhancer, reduced the frequency of expression of *Ly49g*, a natural RME gene, directly tying the enhancer deletion-associated variegation phenomenon to RME. This result powerfully argues that enhancers are not simply permissive for expression at RME genes, but are also instructive regarding expression probability. We hypothesize that the broad range of frequencies with which different *Ly49* genes are naturally expressed (∼5%-60%) in large part reflects differences in enhancer strength. Probabilistic enhancer action has also been documented in *Drosophila*, where regulation of genes by multiple “shadow” enhancers has been suggested to ensure a high probability of gene expression (*40*). We propose that the binary decision to express a gene is regulated by quantitatively varying enhancer activity, which is comprised of both the number of enhancers acting upon a gene and the strength of individual enhancers within that set. Our results are consistent with recent findings that enhancers are probabilistic regulators of transcription burst frequency rather than burst size (*41, 42*). How enhancer control of the probability of stable gene expression interfaces with the control of transcription burst frequency is an exciting area for future investigation.

We predict that the mechanism of mitotic stability of active and silent RME alleles is likely related to maintenance of gene expression states broadly, and may involve bookmarking of promoters (*4*), rather than repressive chromatin at silent alleles. Indeed, we found that expressed and silent *Ly49g* alleles are distinguished only by the presence of active marks at the promoters and gene bodies of active alleles. Enhancers are constitutively accessible and activated on expressed and silent alleles alike, consistent with the decoupling of enhancer and promoter accessibility seen at RME loci generally. The absence of a repressive chromatin state on silent NK receptor alleles suggests that the RME “off state” can reflect a stably inactive, as opposed to repressed, chromatin state. Numerous lineage non-specific genes that are also silent in NK cells (e.g., *Cd19*, *Cd3e*) also lacked traditional repressive chromatin modifications, yet are generally not subject to subsequent activation after an initial failure to be activated. In the case of the NK receptor genes and RME genes broadly it appears that the inactive state is maintained in mature cells in spite of continued enhancer activation, suggesting silent promoters are no longer competent for activation—perhaps due to lack of critical promoter-activating pioneer factor activity that is only present at sufficient levels during differentiation. Notably, whereas some RME genes are expressed in a nonvariegated fashion in progenitor cells (*8, 9*), the NK receptor genes are variegated at the time of initial expression.

Of note, we believe DNA methylation plays at most a minor role since inhibitors of DNA methylation did not appreciably activate expression of silent alleles in our studies or those of others (*43–46*). Significantly, NK receptor gene promoters are CpG poor (*45*). Furthermore, the overwhelming majority of RME genes studied in clones were not responsive to perturbation of DNA methylation (*6, 8, 9*). Our chromatin analyses lead us to propose that binary enhancer action results in two possible outcomes: a stable active state, or a stable inactive state. Functionally, enhancer activity appears to control the probability with which a gene achieves the active state and is therefore classified as lineage-appropriate, while the alternate fate simply resembles inactive lineage non-appropriate genes. This activity varies quantitatively across genetic loci, determining expression likelihood.

The RME of the NK receptor genes resembles the monoallelic expression pattern of cytokine genes including *Il-2, Il-4, Il-5, Il-10* and *Il-13* (*1, 47–49*). The cytokine genes are inducible in response to TCR stimulation and therefore expression is inherently unstable, but impressive stability over several mitotic divisions was observed for *Il-4* (*47*), and the probability of *Il-4* allelic activation and biallelic expression correlated with the strength of the inducing signal (*48*). Intriguingly, the *Il-4* and *Il-13* genes are closely linked and are co-regulated by an enhancer, *CNS1*, that was found to be constitutively acetylated at histone H3, and thus permissive for expression (*50*). Expression of the *Il-4* and *Il-13* gene alleles was independent, however, in a manner strikingly similar to the NK receptor genes. It is probable that the general principles uncovered by us and others apply to genes that are induced via stimulation as well as genes whose expression is acquired during differentiation.

Our data showing RME of *Nkg2d*, *Ptprc*, *CD8a* and *Thy1* suggest that RME is even more prevalent than the previous estimates (*5, 8, 9, 51*). Although RME has been associated previously with poorly expressed genes (*5, 8*), our results extend the phenomenon to relatively highly expressed genes. We propose that genes lie along a spectrum of allelic failure rates that are largely controlled by enhancer strength, with documented RME genes on the highest end of that spectrum. Higher-resolution genome-wide approaches will eventually provide a comprehensive picture of the full extent of RME expression.

Apparently, RME often occurs at such a low rate that it is both beneath ready detection and presumably irrelevant for the function of a cell lineage. In the case of NK receptors, we propose that evolution has exploited RME to generate a complex combinatorial repertoire of NK cell specificities. More speculatively, by regulating expression of fate-determining mediators, RME may underlie stochastic cell fate decisions in some instances of cellular development (*52*). From an evolutionary perspective, appreciable RME of a gene could arise by mutation of strong enhancers of a precursor gene, by providing a new gene with a weak enhancer, or by diminishing the concentration of relevant enhancer-binding transcription factors in a given lineage of cells (as has been shown for several relevant TFs for the *Ly49* genes) (*53, 54*).

Finally, our results suggest that allelic failure rates may in some cases dwarf the rates of null alleles generated by somatic mutation. As a novel mechanism of genetic haploinsufficiency at the cellular level, RME might have broad implications in genetic disease etiology and penetrance of disease phenotypes in heterozygous individuals.

## Acknowledgments

We thank L. Zhang and E. Seidel for assistance, Hector Nola and Alma Valeros in the Cancer Research Laboratory at UC Berkeley for expert assistance with flow cytometry and cell sorting, and M. Wong for assistance with some Illumina sequencing experiments. We thank Drs. Jasper Rine, Russell Vance, Ellen Robey, Michel DuPage, and David Martin for critical evaluation of the manuscript.

This research was supported by NIH grant R01-AI113041 to D.H.R. and in part by the Intramural Research Program of NIH, NIAID (S.A.M).

D.U.K. was supported by a University of California Cancer Research Coordinating Committee Predoctoral Fellowship.

N.K.W. was supported by a National Science Foundation predoctoral fellowship DGE 1752814.

## Author contributions

D.U.K. and D.H.R. conceived and designed the project; D.U.K., N.K.W., C.Z., S.M.D., K.N.M., I.D.P., Y.J. performed cellular and molecular analyses in primary immune cells; S.A.M. provided assistance with sequencing experiments; A.E. and B.C. provided bioinformatic support; D.U.K. and D.H.R. wrote the manuscript incorporating feedback from all authors.

## Data and materials availability

ATAC-seq and CUT&RUN datasets are available as an NCBI Gene Expression Omnibus (GEO) series under the accession number GSE181197. Prior to publication, a secure token is required to access this data series.

## Materials and Methods

### Animals and animal procedures

All mice were maintained at the University of California, Berkeley. *Nkg2d^-/-^* mice are available at the Jackson Laboratory (JAX Stock No. 022733). C57BL/6J (B6), BALBc/J and B6;129-*Ncr1^tm1Oman^*/J (Ncr^gfp^) were purchased from the Jackson Laboratory and bred at UC Berkeley. BALB/cJ, CBA/J and AKR/J and B6.SJL-*Ptprc^a^ Pepc^b^*/BoyJ mice were purchased from the Jackson Laboratory. F_1_ hybrid mice were purchased from the Jackson Laboratory or were generated at UC Berkeley from inbred parents.

For the generation of CRISPR edited mice, Cas9 RNP was delivered to single-cell embryos either through microinjection or CRISPR-EZ electroporation, both of which are described in reference (*57*). *Ly49a_Hss1Δ_* mice were generated by microinjection, while *Nkg2a_5’EΔ_*, *Nkg2d_5’EΔ_* and *Ly49g_5’EΔ_* mice were generated by CRISPR-EZ electroporation. Whether through microinjection or electroporation, we used paired sgRNAs flanking the enhancer to generate enhancer deletion mice. sgRNAs were selected using the GPP web portal from the Broad Institute. Guides with highest predicted editing efficiencies were prioritized, while also minimizing for predicted off-target cutting in protein-coding genes. sgRNAs were generated using the HiScribe T7 Quick High Yield RNA Synthesis Kit (New England Biolabs). Founder mice (F_0_) harboring deletion alleles were backcrossed to C57BL/6J (B6) mice to generate heterozygous F_1_ mice, and were then intercrossed to generate WT, heterozygous and homozygous littermates for experiments. All sgRNAs used for the generation of enhancer deletion mice are listed in Table S1. Primers used to PCR identify edited founders and genotype subsequent filial generations are listed in Table S1. All animals were used between 8-32 weeks of age, and all experiments were approved by the UC Berkeley Animal Care and Use Committee (ACUC).

### Flow Cytometry

Single cell splenocyte suspensions were generated by passing spleens through a 40 μm filter. Red blood cells were lysed with ACK buffer. Fresh splenocytes, or where indicated cells cultured with 1000 U/ml recombinant human IL-2 (National Cancer Institute) were stained for flow cytometry in PBS containing 2.5% FCS (FACS Buffer). Before staining with antibodies, Fc*γ*RII/III receptors were blocked for 15 minutes at 4C using 2.4G2 hybridoma supernatant. Cells were washed with FACS buffer and then stained with antibodies directly conjugated to fluorochromes or biotin at 4°C for 15 to 30 minutes. In order to differentiate between alleles of a receptor in (B6 x BALB/c)F_1_ hybrid NK cells, the B6-specfic clone was used first in order to block epitopes in competition with the clone recognizing both alleles. For example, to discriminate Ly49G2 alleles, cells were stained for at least 15 minutes with 3/25 which recognizes Ly49G2^B6^, and then 4D11 was added. For discriminating alleles of NKG2A, cells were stained first with the NKG2A^B6^-specific 16al1, followed by 20d5, which binds to both alleles. Ly49A^B6^ (A1) was added before the non-discriminating JR9 clone, but in this case, cells expressing only the B6 allele did not resolve from the population of cells expressing both alleles. When necessary, cells were washed and then stained with secondary antibody or fluorochrome-conjugated streptavidin. Near-IR viability dye (Invitrogen L34975) or DAPI (Biolegend 422801) were used to discriminate live cells. Flow cytometry was carried out using an LSR Fortessa or X20 from BD Biosciences, and data were analyzed using FlowJo software. In all cases, NK cells were defined as CD3^-^NKp46^+^ splenocytes. For sorting on a BD FACSAria II sorter, the samples were prepared nearly identically as they were for flow cytometric analysis with the exception that the medium used was sterile RPMI 1640 (ThermoFisher) with 5% FCS.

### Antibodies used in flow cytometry

From Biolegend: anti-CD3*ε* (145-2C11) PE-Cy5, anti-CD4 (GK1.5) BUV737, anti-CD19 (6D5) PE-Cy5, anti-F4/80 (BM8) PE-Cy5, anti-Ter119 (TER-119) PE-Cy5, anti-NKp46 (29A1.4) BV421, anti-NKG2A^B6^ (16a11) PE, anti-Ly49A^B6^ (A1) PE, anti-NKG2D (CX5) PE/Dazzle 594, anti-CD8*β* (YTS156.7.7) PE-Cy7, anti-CD45.1 (A40) APC, anti-CD45.2 (104) FITC, anti-CD90.2 (53-2.1) PE or FITC, goat-anti-mouse IgG (Poly4053) PE. From eBioscience/ThermoFisher: anti-NKG2A (20d5) PerCP, anti-Ly49I (YLI-90) FITC, anti-Ly49G2 (4D11) PerCP or PE-Cy7, anti-CD90.1 (HIS51) FITC, anti-rat IgG F(ab’)2 (polyclonal, lot 17-4822-820) APC. From BioXCell: anti-CD8.1 (116-13.1) unconjugated primary mouse IgG2a *κ*, anti-CD8.2 (2.43) unconjugated primary rat IgG. Purified in-house: anti-Ly49A (JR9) biotin/APC streptavidin from Biolegend, anti-Ly49G2^B6^ (3/25), anti-NKG2D (MI6) biotin/PE Streptavidin from Biolegend.

### Ex vivo NK cell cultures

NK cells were prepared from spleens by passage through a 40 μm filter. Red blood cells were lysed with ACK. Splenocytes were cultured in RPMI 1640 (ThermoFisher) with 1000 U/mL IL-2 (National Cancer Institute) and 5% FCS. In all cases, media was supplemented with 0.2 mg/mL glutamine (Sigma), 100 U/mL penicillin (ThermoFisher), 100 μg/mL streptomycin (Thermo Fisher Scientific), 10 μg/mL gentamycin sulfate (Fisher Scientific), 50 μM β-mercaptoethanol (EMD Biosciences), and 20 mM HEPES (ThermoFisher).

### Analysis of the stability of monoallelic expression of NKG2D

NKG2D+/-NK cells were sorted from *WT* or *Nkg2d^5’EΔ/5’EΔ^* mice on day 2 or 3 of *ex vivo* NK cell culture in IL-2 medium as described above. Cells were cultured *in vitro* in IL-2 containing media for a further 8-10 days, during which cells expanded ∼10-100 fold based on hemocytometer counts. Cells were analyzed for NKG2D expression by flow cytometry. In all cases medium contained 5% FCS (Omega Scientific), 0.2 mg/mL glutamine (Sigma), 100 U/mL penicillin (ThermoFisher), 100 μg/mL streptomycin (Thermo Fisher Scientific), 10 μg/mL gentamycin sulfate (Fisher Scientific), 50 μM β-mercaptoethanol (EMD Biosciences), and 20 mM HEPES (ThermoFisher). Cells were incubated at 37°C in 5% CO_2_.

### Ex vivo assay for the stability of monoallelic expression in T cells

Cells from the spleens and a collection of lymph nodes (brachial, axial, inguinal, mesenteric) from F_1_ hybrid mice and parental inbred line controls were combined and passed through a 40 μm filter, and red blood cells were lysed with ACK buffer. Cells were prepared for sorting as described above, staining with the relevant allele-specific antibodies. For CD45 monoallelic expression, Thy1+ cells were further gated according to CD45 allelic expression. For CD8*α* monoallelic expression, CD3+CD8*β*+ cells were analyzed for CD8*α*allelic expression. For Thy1 monoallelic expression, CD4+MHC II-cells were analyzed for Thy1 allelic expression. Cells expressing either the paternal or maternal allele (or both) of the receptor studied were sorted and expanded for 1 week in RPMI 1640 (ThermoFisher) containing 200 U/mL recombinant IL-2, Dynabeads mouse T-activator CD3/CD28 (ThermoFisher) beads at a 1:1 cells to beads ratio, 10% FCS, and supplemented as RPMI 1640 above. After 1 week of expansion, cells were harvested, counted by hemocytometer and prepared for a second sort. After sorting for expression of the relevant receptor allele again in order to ensure purity, cells were once again expanded in a restimulation, this time with a cells to beads ratio of 10:1. After the second expansion, cells were again counted, stained and prepped for final analysis of monoallelic receptor expression by flow cytometry.

In analysis of *Ptprc* monoallelic expression, RNA was isolated from expanded T cells expression either or both *Ptprc* alleles as displayed in Fig. 7D using the iScript cDNA synthesis kit (BioRad), from 10,000-40,000 cells. Half of the reaction volume (10μL out of 20μL) were used to PCR amplify a region of the *Ptprc* transcript using intron-spanning PCR primers (Table S3).

### Enhancer deletion in primary NK cells via Cas9-RNP nucleofection

*Ex vivo* editing of primary mouse NK cells was carried out according to a modified version of the protocol used to modify primary human T cells described in reference (*34*). Cas9 was purchased from the UC Berkeley Macro Lab core (40 uM Cas9 in 20 mM HEPES-KOH, pH 7.5, 150 mM KCl, 10% glycerol, 1 mM DTT), and sgRNAs were transcribed *in vitro* according to the Corn lab online protocol (https://www.protocols.io/view/in-vitro-transcription-of-guide-rnas-and-5-triphos-bqjbmuin). NK cells were prepared by sorting day 5 IL-2 cultured NK cells from (B6 x BALB/c)F_1_ hybrids. CD3-NKp46+ Cells were sorted to be positive for either NKG2A^B6^ using the 16a11 clone or Ly49G2^B6^ using the 3/25 clone, and cells were further cultured overnight in RPMI 1640 media containing 5% FCS and 1000 U/mL IL-2 (National Cancer Institute). On day 6, 1 million sorted NK cells were prepared for nucleofection using the Lonza 4D-Nucleofector per condition. Cas9 and sgRNAs were complexed at a molar ratio of 1:2 (2.5 μL of 40 μM Cas9 was added to 2.5 μL of sgRNA suspended at 80 μM (6.5 μg) in nuclease-free H_2_O). If two flanking guides were used, 1.25 μL of each were used, maintaining the Cas9 to sgRNA molar ratio. Cas9-RNP was complexed for 15 minutes at 37°C and transferred to a single well of a 96-well strip nucleofection cuvette from Lonza for use with the Nucleofector 4D. 1 million sorted day 6 IL-2 cultured NK cells were resuspended in 18 μL of supplemented Lonza P3 buffer from the P3 Primary Cell kit, and added to the Cas9-RNP complex. Cells were nucleofected using the CM137 nucleofection protocol and 80 μL pre-warmed RPMI 1640 with 5% FCS was immediately added. After a 15-minute recovery period at 37°C, cells were returned to culture in 1 mL of RPMI 1640 with 5% FCS and 1000 U/mL IL-2. After 5-7 days in culture maintaining a density of approximately 1 million cells/mL, receptor expression was assayed by flow cytometry. In order to validate enhancer flanking guides (Table S1) an identical protocol was followed with either day 5 IL-2 cultured splenocytes, or day 5 IL-2 cultured NK cells isolated using the MojoSort NK isolation kit from Biolegend, but instead of analysis by flow cytometry, gDNA was prepared and used as a template for PCR to detect the expected deletion.

### F_1_ hybrid genetics and calculations of expected changes in receptor-expressing NK cell populations

F_1_ hybrid genetics were carried out by breeding WT or CRISPR/Cas9-edited males on the B6 background to females from the following backgrounds: BALBc/J, CBA/J, AKR/J. Edited alleles were crossed only to BALBc/J, while CBA/J and AKR/J were used in the F_1_ hybrid analysis of monoallelic expression of CD8*α* and Thy1, respectively.

We estimated the expected frequencies of NK cells in (*Nkg2a^B6-5’E^*^Δ^*^/BALB/c+^*) F_1_ mice by assuming independence of allelic expression. That assumption leads to the following predictions:

The percentage of cells expressing neither allele in the mutant will equal the sum of the percentages of the two NK cell populations that lack NKG2A^BALB/c^ in WT (B6 x BALB/c)F_1_ hybrids, that is the cells that express neither allele, and cells expressing only the B6 allele.
The percentage of cells expressing NKG2A^BALB/c^ only in the mutant will equal the sum of the percentages of the NK cell populations in WT (B6 x BALB/c)F_1_ hybrids that express NKG2A^BALB/c^, that is the cells that express only the BALB/c allele, and the cells expressing both alleles.
The percentages of cells expressing NKG2A^B6^ only or both NKG2A^B6^ and NKG2A^BALB/c^ will be 0, since NKG2A^B6^ is not expressed,

The expected frequency of cells expressing NKG2D in *Nkg2d^-/5’E^*^Δ^ mice was calculated assuming stochastic expression of alleles, and was based on the frequency of cells expressing NKG2D, or not, in *Nkg2d^5’E^*^Δ*/5’E*Δ^ mice. The frequency of cells lacking expression of a given allele is the square root of the frequency of cells expressing neither allele. Subtraction of this proportion from 1 yields the predicted frequency of cells expressing NKG2D in *Nkg2d^-/5’E^*^Δ^ mice. E.g., an observed NKG2D expression frequency of ∼67% in a *Nkg2d^5’E^*^Δ*/5’E*Δ^ mouse would result in an expected frequency datapoint of ∼43%.

The expected changes in populations with respect to Ly49G2 alleles in *Ly49g^B6-Hss5^*^Δ/*BALB/c+*^ mice were calculated with the same assumption of independent regulation of alleles.

We started by calculating the overall percentage of cells expressing Ly49G2^B6^ in the F1 mice with the mutation, which averaged 47.7% of that in WT F1 mice.
The predicted percentage of cells expressing only Ly49A^B6^ in the mutant F1 was then 47.7% of the percentage of cells expressing only Ly49A^B6^ in WT mice.
And the predicted percentage of cells expressing both alleles in the mutant F1 was 47.7% of the percentage of cells expressing both alleles in WT mice.
The predicted percentage of cells expressing neither allele in the mutant F1 was calculated as the percentage of cells expressing neither allele in WT mice + 52.3% (100%-47.7%) of the precentage of NK cells that express only Ly49G2^B6^ in WT mice.
Finally, the predicted percentage of NK cells expressing only Ly49G2^BALB/c^ in the mutant was calculated as the percentage expressing only Ly49G2^BALB/c^ in WT mice plus 52.3% of the NK cells expressing both alleles in WT mice.

Note that the genetic background of the mice significantly influences *Ly49g* expression even in WT mice, presumably reflecting trans-acting events (e.g. each *Ly49g^B6+^* allele is expressed on ∼31% of NK cells in B6 mice, but only ∼19% in F_1_ hybrid mice). Therefore expected data are calculated using Ly49G2^B6^ expression frequencies in *Ly49g^B6-5’E^*^+/*BALB/c+*^ mice.

### ATAC-seq

ATAC-seq was performed as previously described in reference (*58*). Briefly, 50,000 sorted NK cells were washed in cold PBS and resuspended in lysis buffer (10 mM Tris-HCl, pH 7.4; 10 mM NaCl; 3 mM MgCl_2_; 0.1% (v/v) Igepal CA-630). The crude nuclear prep was then centrifuged and resuspended in 1x TD buffer containing the Tn5 transposase (Illumina FC-121-1030). The transposition reaction was incubated at 37C for 30 minutes and immediately purified using the Qiagen MinElute kit. Libraries were PCR amplified using the Nextera complementary primers listed in reference (*58*) and were sequenced using an Illumina NextSeq 500 or a HiSeq 4000.

### CUT&RUN

CUT&RUN was performed essentially as previously described (*59*). Briefly, 50,000-500,000 NK cells were washed and immobilized on Con A beads (Bangs Laboratories) and permeabilized with wash buffer containing 0.05% w/v Digitonin (Sigma-Aldrich). Cells were incubated rotating for 2 hours at 4°C with antibody at a concentration of 10-20 μg/mL. Permeabilized cells were washed and incubated rotating at room temperature for 10 minutes with pA-MNase (kindly provided by the Henikoff lab) at a concentration of 700 ng/mL. After washing, cells were incubated at 0°C and MNase digestion was initiated by addition of CaCl_2_ to 1.3 mM. After 30 minutes, the reaction was stopped by the addition of EDTA and EGTA. Chromatin fragments were released by incubation at 37°C for 10 minutes, purified by overnight proteinase K digestion at a concentration of 120 μg/mL with 0.1% wt/vol SDS at 55°C. DNA was finally purified by phenol/chloroform extraction followed by PEG-8000 precipitation (final concentration of 15% wt/vol) using Sera-mag SpeedBeads (Fisher) (https://ethanomics.files.wordpress.com/2012/08/serapure_v2-2.pdf).

Libraries were prepared using the New England Biolabs Ultra II DNA library prep kit for Illumina as described online (https://www.protocols.io/view/library-prep-for-cut-amp-run-with-nebnext-ultra-ii-bagaibse?version_warning=no) with the following specifications and modifications. The entire preparation of purified CUT&RUN fragments from a reaction were used to create libraries. For histone modifications, end repair and dA-tailing were carried out at 65°C. NEB hairpin adapters (From NEBNext Multiplex Oligos for Illumina) were diluted 25-fold in TBS buffer and ligated at 20°C for 15 minutes, and hairpins were cleaved by the addition of USER enzyme. Size selection was performed with AmpureXP beads (Agencourt), adding 0.4X volumes to remove large fragments. The supernatant was recovered, and a further 0.6X volumes of AmpureXP beads were added along with 0.6X volumes of PEG-8000 (20% wt/vol PEG-8000, 2.5 M NaCl) for quantitative recovery of smaller fragments. Adapter-ligated libraries were amplified for 15 cycles using NEBNext Ultra II Q5 Master Mix using the universal primer and an indexing primer provided with the NEBNext oligos. Amplified libraries were further purified with the addition of 1.0X volumes of AmpureXP beads to remove adapter dimer and eluted in 25 μL H_2_O. Libraries were quantified by Qubit (ThermoFisher) and Bioanalyzer (Agilent) before sequencing on an Illumina HiSeq 4000 or MiniSeq as paired-ends to a depth of 10-32 million.

The following antibodies were used for CUT&RUN: Abcam: anti-H3K4me1(ab8895), anti-H3K4me2 (ab7766), anti-H3K4me3 (ab8580), anti-H3K27ac (ab4729), anti-H3K9me3 (ab8898). Cell Signaling: anti-H3K27me3 (C36B11), anti-H2AUb1 (D27C4). Control IgG (cIgG) from Biolegend: Mouse IgG2a*κ* (MOPC-173)

### Datasets and processing and visualization

Raw mined datasets were downloaded from NCBI Gene Expression Omnibus (GEO) or the European Bioinformatics Institute (EBI). NK cell ATAC-seq and histone modification (H3K4me1, H3K4me2, H3K4me3, H3K27ac) were from reference (*55*) under GEO accession numbers GSE59992 and GSE60103. Runx3 ChIP-seq data and non-immune serum control in NK cells were sourced from reference (*60*) (GSE52625) and T-bet ChIP-seq data and input control were sourced from reference (*61*) (GSE77695). p300 ChIP-seq raw data was sourced from reference (*56*) (GSE145299). p300 ChIP-seq peaks were called in reference (*56*) and downloaded in .csv format.

Raw data from all datasets (mined or generated in this study) were processed using an in-house assembled pipeline. Datasets were tested with FastQC. Paired-end reads were then aligned to the mm10 reference genome using Bowtie2 with the --sensitive parameter. Paired-end CUT&RUN libraries were tested and aligned with the same pipeline. All reads aligned to the mitochondrial chromosome were removed with samtools. Aligned reads were then sorted, indexed, and filtered for a mapping quality of ≥10 with samtools. PCR duplicates were removed with Picard (Broad Institute). Reads covering blacklisted regions (ENCODE mm10 database), were removed with bedtools. Data were then normalized to signal per million reads (SPMR) when calling narrow peaks with macs2. Resultant bedgraph files were converted to bigwigs with the bedGraphToBigWig program from the UCSC Genome Browser toolkit for visualization on Integrative Genomics Viewer (IGV) (*62*). Data in Fig. S1A were plotted using the Bioconductor package SeqPlots (*63*).

### Ranking of accessible sites in NK cells according to H3K4me1:me3 ratio

Reads from duplicate ChIP-seq datasets (for both H3K4me1 and H3K4me3) from reference (*55*) were merged to ensure robust signal, and the resultant files were processed and normalized as above. NK cell ATAC-seq peaks were called in the Ly49G2^B6+BALB+^ NK cell ATAC-seq dataset using macs2 narrowpeaks. Before ranking, ATAC-seq peaks were filtered such that only peaks that fell within the top 95% of both H3K4me1 and H3K4me3 signal computed over a 2 kb window from the peak midpoint computed using pandas and numpy in Python 3.7.4, resulting in 51,650 usable peaks. H3K4me1:me3 raw ratio and log2 ratio bigwigs were generated with the bamCompare utility from deepTools (v2.5.4). The log2 ratio track was visualized on IGV, and the raw ratio was used to rank ATAC-seq peaks. Heatmaps were generated with the computeMatrix and plotHeatmap utilities from deepTools (v2.5.4). Heatmaps were sorted by the mean H3K4me1:me3 ratio signal over a 2 kb window centered at the midpoint of the 51,650 ATAC-seq peaks. *Hss1* and *5’E* enhancer regions and corresponding promoters at NKC genes were individually predefined and the position of each was then marked on the heatmap.

### Definition of NK cell promoters and enhancers and ranking of regulatory elements according to H3K4me1:me3 ratio

Annotated mouse promoters (defined as the TSS at a single nucleotide) in the mm10 genome assembly were downloaded as a BED file from the EDPNew database (*33*). To identify likely active promoters in NK cells, broad regions of H3K27ac were called based on ChIP-seq data sourced from reference (*55*) using the “macs2 callpeak --broad” command. Mouse EDPNew promoters falling within broad H3K27ac domains were identified using the “bedtools intersect - wa” command, resulting in a set of 9901 active promoters in mouse NK cells.

Enhancers in naïve mouse NK cells were defined as the intersection of ATAC-seq and p300 peaks not found at the promoters as defined above. p300 ChIP-seq peaks in resting NK cells were previously defined and downloaded from reference (*56*). ATAC-seq peaks that were enriched in p300 binding were identified using the “bedtools intersect -wa” command. To define enhancers that do not overlap annotated promoters, EDPNew promoters were subtracted from p300-enriched ATAC-seq peaks using the “bedtools subtract” command resulting in 10,246 NK cell enhancers.

### SNPsplit chromosome of origin reads analysis

Delineation of allele-informative reads was performed similarly as in reference (*4*). SNPs between the C57BL/6 (B6) and BALB/cJ (BALB) mouse strains were sourced from the Wellcome Sanger Institute Mouse Genomes Project dbSNP (v142). In order to perform unbiased alignment of reads originating from both the B6 and BALB genomes, SNPs marked by the database were replaced by ‘N’ in the mm10 reference genome that we use for alignment using SNPsplit (Babraham Institute) (*36*). ATAC-seq datasets generated in (B6 x BALB) F_1_ hybrid NK cells were then aligned to the N-masked genome using bowtie2 and further processed and normalized as above. Reads that overlapped the annotated SNPs were marked as allelically informative reads after alignment and quality control using SNPsplit. Allele-informative reads were then processed and normalized as described above. ∼4% of ATAC-seq reads across the dataset were allele-informative.

### ChromHMM construction of 3 state model

CUT&RUN data for four histone modifications (H3K4me3, H3K9me3, H3K27ac, H2AK119ub1) generated in cells expressing neither allele (DN) or both alleles (DP) of Ly49G2 were separately used to construct chromatin states using ChromHMM (v1.22) (*39*). The genome was segmented into three distinct states: state 1 (active chromatin; enriched in H3K4me3 and H3K27ac), state 2 (inactive chromatin; lacking enrichment of all four marks), and state 3 (repressed chromatin enriched in H3K9me3 and H2AK119ub1). The resultant .bed file outputs were visualized with IGV.

### Statistical analysis

*In vivo* germline-edited mouse data were compared with One-way ANOVAs with Tukey’s multiple comparisons (when three or more genotypes were compared) or student’s *t-*tests (when only two groups were compared). *Ex vivo* edited NK cell experiments were analyzed by ratio paired *t*-tests comparing experimental and control samples within a single experiment. In all cases, \**P* <0.05; \*\**P* <0.01; \*\*\**P* <0.001; \*\*\*\**P* <0.0001.

---

**Fig. S1.**
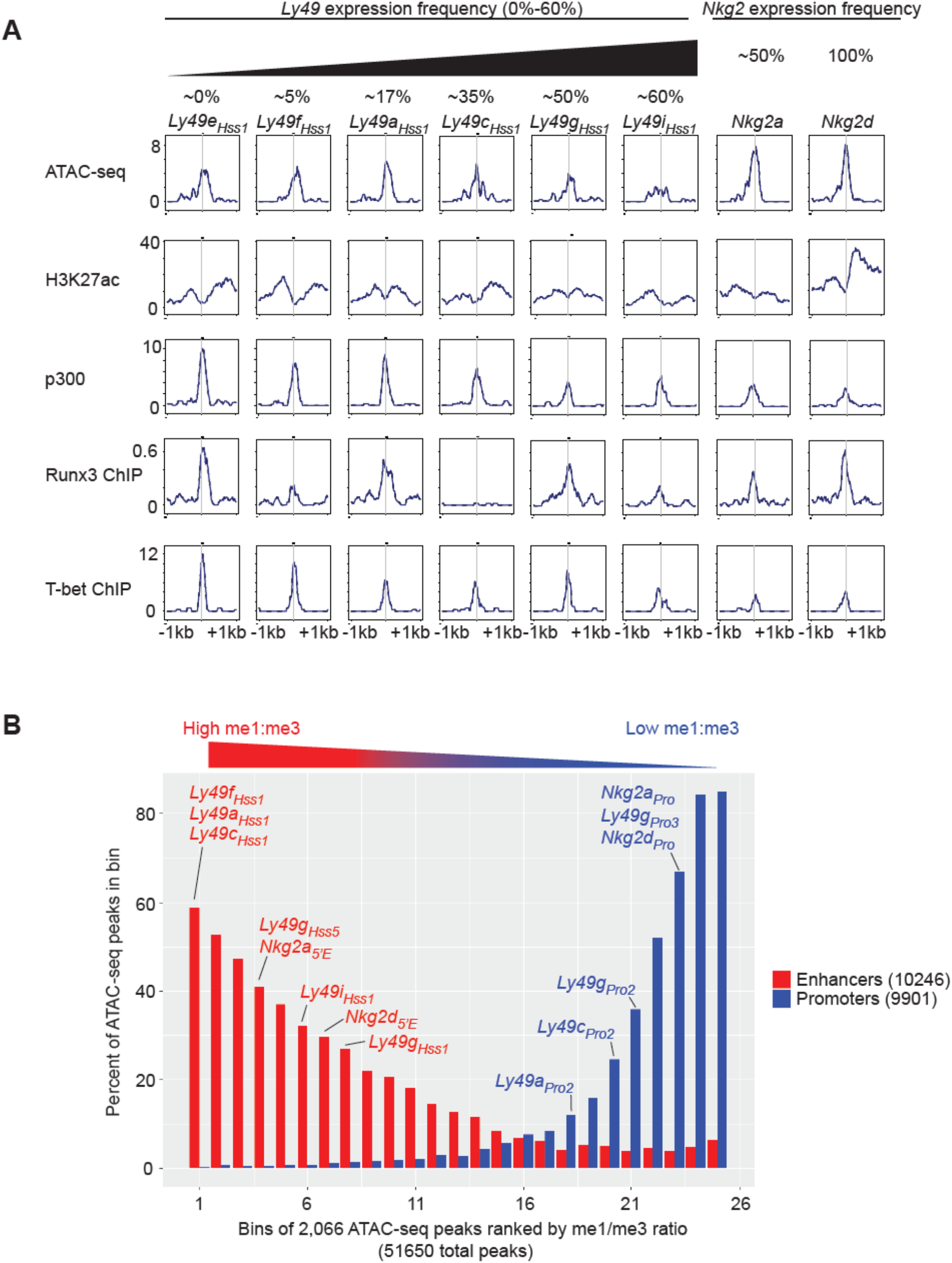
Chromatin features and TF binding profile of the *Ly49_Hss1_* and *Nkg2_5’E_* enhancers. **(A)** Selected *Ly49* genes are depicted left to right according to expression percentages, which are specified above each gene name. *Nkg2_5’E_* elements are depicted on the right. ATAC-seq and ChIP-seq data profiles are shown over a 2 kb window centered at the midpoint of the called ATAC-seq peak. ATAC-seq and H3K27ac are sourced from ref (*55*), p300 data are from ref (*56*), Runx3 data are from ref (*60*) and T-bet data are from ref (*61*). **(B)** The H3K4me1 and H3K4me3 ChIP-seq datasets analyzed in Fig. 1 from ref (*55*) were used to examine these modifications in 51,560 MACS2-called ATAC-seq peaks in NK cells expressing both Ly49G2 alleles (Fig. 2D). The MACS2-called peaks were first filtered for peaks found in the top 95% of both me1 and me3 signal in NK cells, and then ranked by me1:me3 ratio over a 2kb window as in Fig. 1C. The filtered ATAC-seq peaks were then binned in sets of 2,066 peaks according to me1:me3 ratio, with the highest me1:me3 ratio as bin 1. Separately, a total of 9,901 NK cell promoters were defined by mouse EPDnew as promoters that overlap with broad H3K27ac peaks called from sourced from ref (*55*), and a total of 10,246 enhancers were defined as ATAC-seq peaks that are enriched in p300 ChIP-seq signal, sourced from ref (*56*). Within each bin, the percentage of peaks that overlap with enhancers (red) or promoters (blue) defined in this manner are depicted. Bins that contain ATAC-seq peaks corresponding to key selected NK receptor gene promoters or enhancers are indicated.

**Fig. S2.**
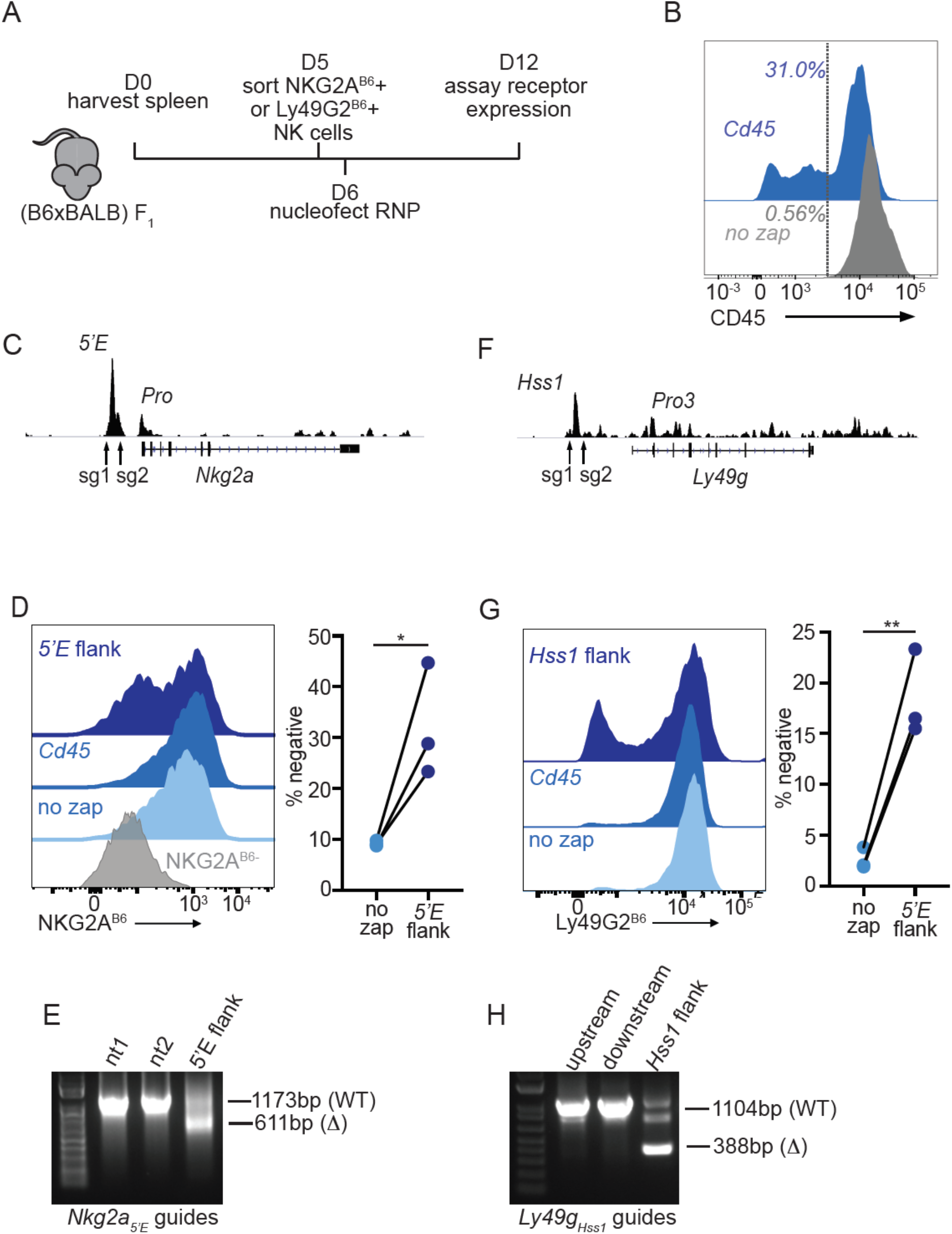
The *Nkg2a_5’E_* and *Ly49g_Hss1_* enhancers are required to maintain gene expression in primary NK cells. (**A**) Experimental design. (B6 x BALB/c)F_1_ splenocytes were cultured with IL-2 for 5 days before sorting NK cells positive for NKG2A^B6^ or Ly49G2^B6^. After recovery, cells were nucleofected with Cas9-RNP complexed with the indicated sgRNA, or were not treated (“no zap”) before culture and analysis. (**B**) CD45 staining of IL-2 cultured NK cells isolated from splenocytes using the Mojosort kit on day 2 of culture, nucleofected, or not, with an sgRNA to disrupt the *Cd45* gene, on day 5 of culture and stained for CD45 expression on day 10 of culture. (**C**) Location of flanking guide RNAs for deleting *Nkg2a_5’E_* with NK cell ATAC-seq data for reference. (**D**) Flow cytometric analysis of NKG2A^B6^ expression by NK cells on day 12 (6 days after nucleofection with sgRNAs (5’E flank), control CD45 sgRNAs or no treatment). Control cells were sorted to be NKG2A^B6^-negative on day 6 (grey). Data from 3 independent experiments are quantified on the right. (**E**) Deletion test of sgRNAs used to delete *Nkg2a_5’E_*. Agarose gel electrophoresis of PCR amplified fragments from the region surrounding *Nkg2a_5’E_* in cells nucleofected with either non-targeting (nt) guides, or guides flanking *NKG2a_5’E_* (5’E flank) in IL-2 cultured mouse splenocytes. Amplicon sizes of the WT and D bands are depicted. NK cells enriched using the MojoSort Mouse NK cell isolation kit from Biolegend were cultured for 6 days in IL-2 containing media before samples of 1 x 10^6^ cells were nucleofected on day 6. On day 9, gDNA was prepared and used as a template for PCR. (**F**-**G**) Flow cytometric analysis of Ly49G2^B6^ expression in cells nucleofected with *Ly49g_Hss1_* sgRNAs, control sgRNAs or no treatment, as in (C-D). Data from 3 independent experiments are quantified on the right. The groups were compared using ratio paired *t*-tests. \**P < 0.05*, \*\**P < 0.01*. (**H**) Deletion test, as in (E) for the sgRNA pair used to delete *Ly49g_Hss1_* in comparison to results with only the upstream or downstream gRNAs.

**Fig. S3.**
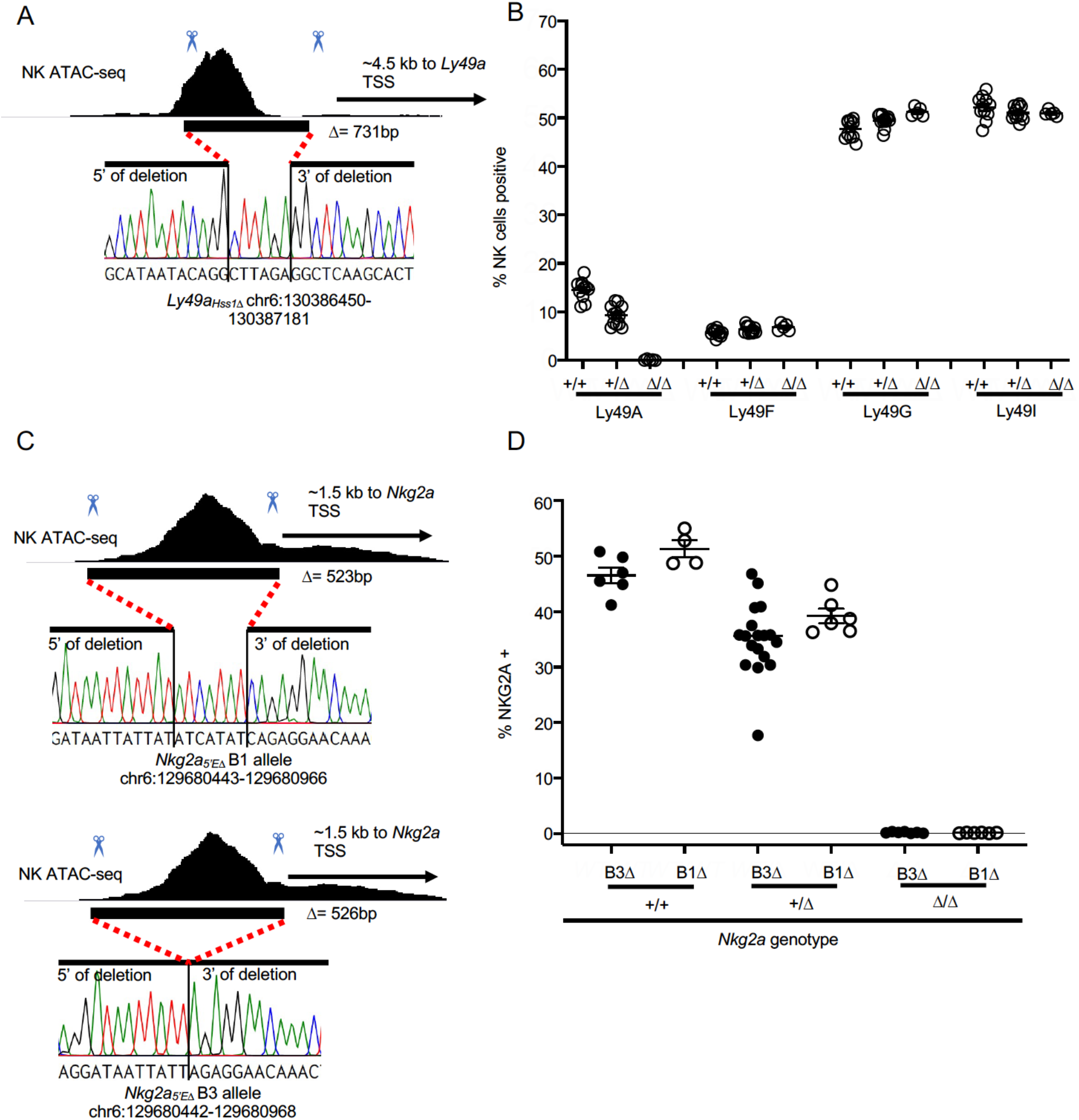
*Ly49a_Hss1_*_Δ_ and *Nkg2a_5’E_*_Δ_ alleles employed in the study. (**A**) Genomic position and sequence of the *Ly49a_Hss1_*_Δ_ allele analyzed in this study. The black bar shows the location of the CRISPR/Cas9 generated in/del based on Sanger sequencing of a PCR amplicon spanning the region. (**B**) Percentages of cells expressing indicated Ly49 receptors in *Ly49a^Hss1^*^Δ/*Hss1*Δ^ mice, heterozygous and wildtype littermates, from flow cytometry analyses. Data are combined from two independent experiments, n=5-12. (**C**) Two *Nkg2a_5’E_*_Δ_ alleles generated and analyzed in this study, as in panel (A). (**D**) Percentages of cells expressing NKG2A in mice with the genotypes shown. Data are combined from two independent experiments with the *Nkg2a_5’E_*-B3Δ allele, and one experiment with the *Nkg2a_5’E_*-B1Δ allele.

**Fig. S4.**
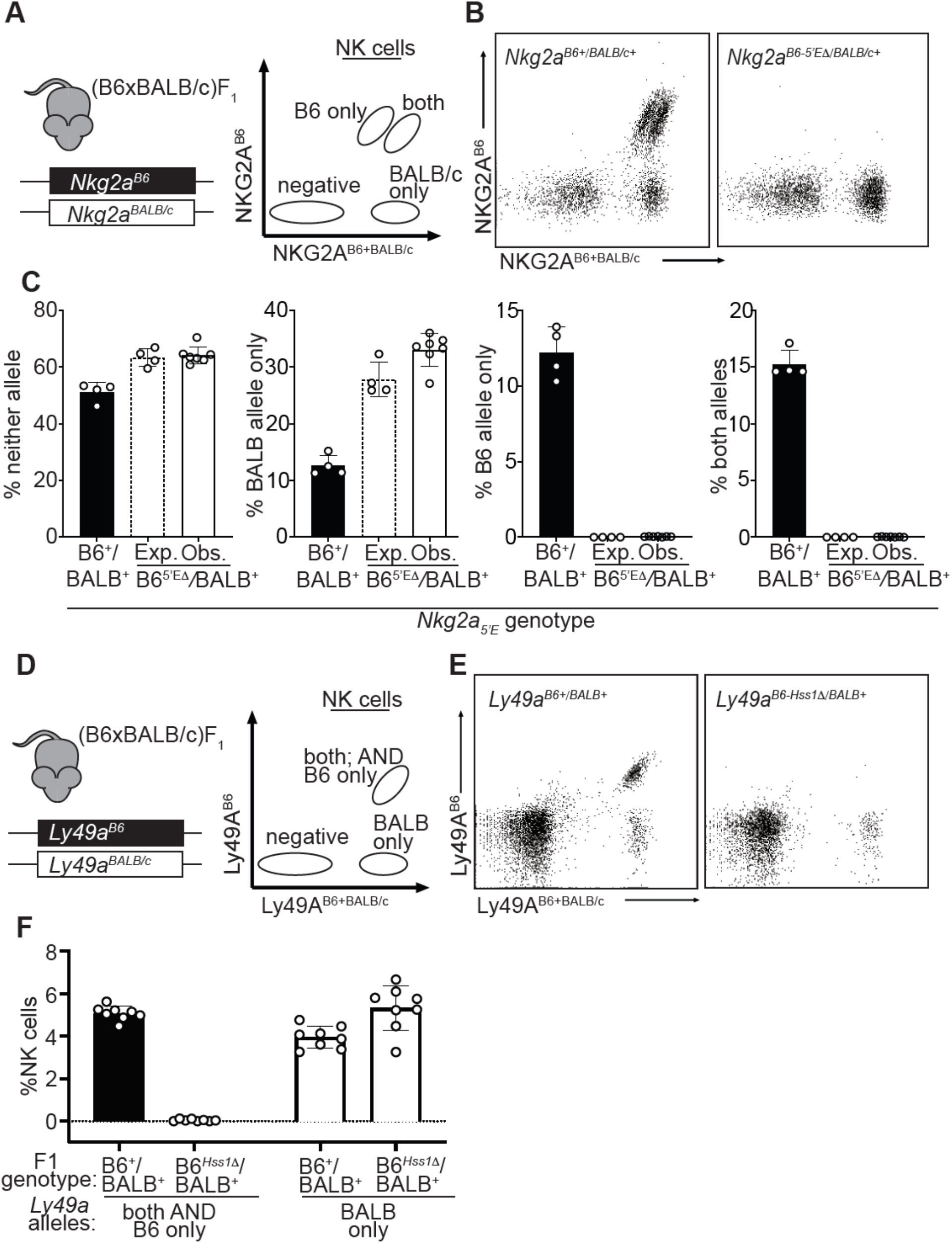
The constitutively accessible *Nkg2a_5’E_* and *Ly49a_Hss1_* enhancers act entirely *in cis*. (**A**) Schematic of (B6 x BALB/c)F_1_ hybrid NK cell staining pattern using 16a11 (NKG2A^B6^ reactive) and 20d5 (NKG2A^B6+BALB/c^ reactive) antibodies. (**B**) Representative dot plots displaying staining of (B6 x BALB/c)F1 hybrid splenic NK cells using 16a11 and 20d5 (*n*=4-7). (**C**) Expected (dotted bar) and observed (solid white bar) percentages of populations in *Nkg2a^B6-5’E^*^Δ^*^/BALB/c-5’E+^* mice, compared to wildtype littermate (B6 x BALB/c)F1 hybrid mice (black bar). Expected frequencies are calculated assuming stochastic *cis* regulation of alleles (detailed in methods). Data are representative of two independent experiments. (**D**) Schematic of (B6 x BALB/c)F_1_ NK cell staining pattern using A1 (Ly49A^B6^ reactive) and JR9 (Ly49A^B6+BALB/c^ reactive) antibodies. (**E**) Representative dot plots displaying (B6 x BALB/c)F1 hybrid NK cells using A1 and JR9. (**F**) Percentages of NK cells expressing the indicated Ly49A alleles; data are combined from two independent experiments (*n*=8-9). Error bars in all panels denote SEM.

**Fig. S5.**
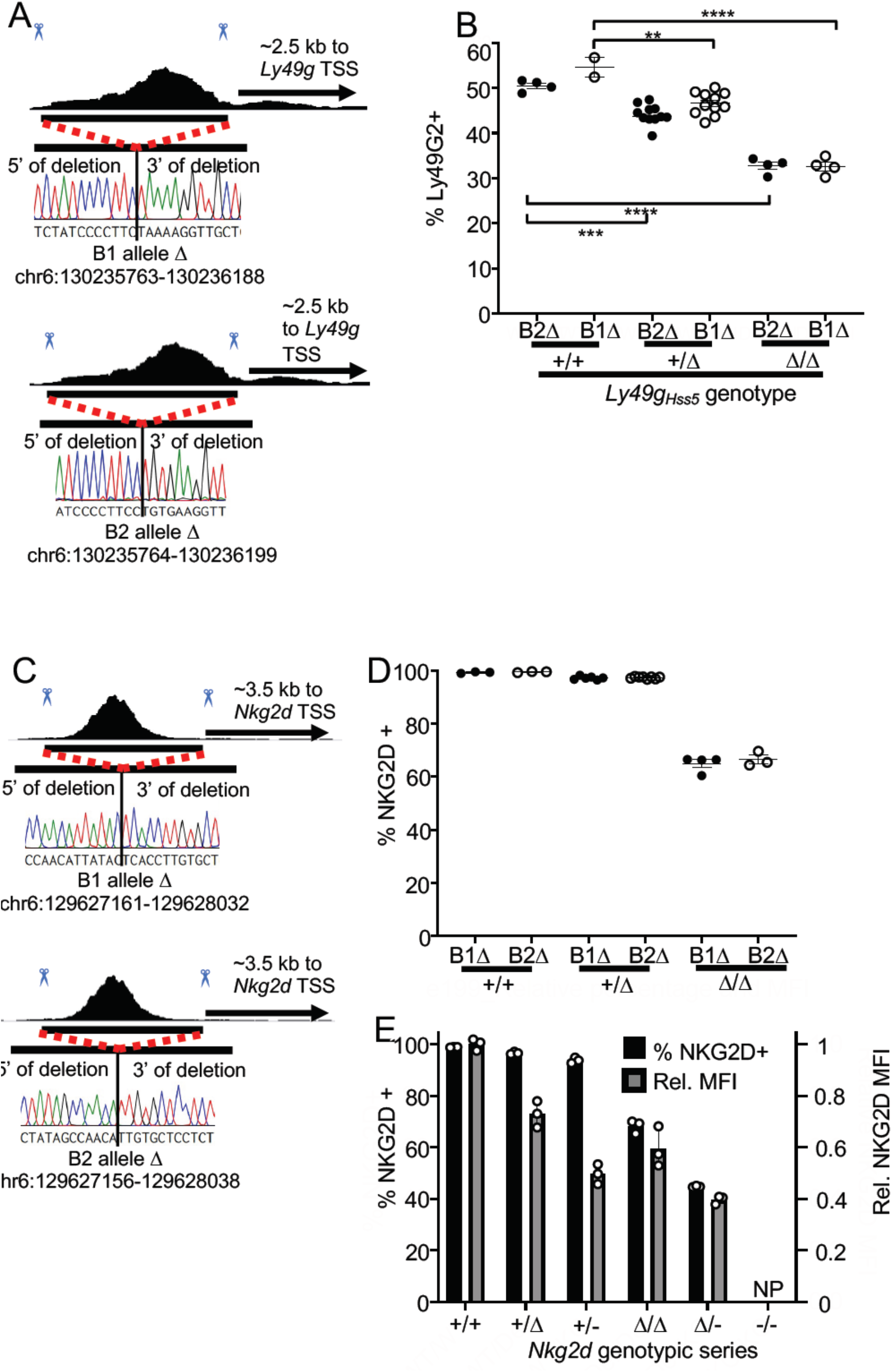
*Ly49g_Hss5_*_Δ_ and *Nkg2d_5’E_*_Δ_ alleles employed in this study. (**A**) Genomic position and sequences of *Ly49g_Hss5_*_Δ_ alleles as in fig. S3A. (**B**) Percentages of cells expressing Ly49G2 in mice with the indicated genotypes, comparing one experiment each with the two alleles (*Ly49g_Hss5_* -B1Δ and *Ly49g_Hss5_*-B2Δ). \*\**P* <0.01; \*\*\**P* <0.001; \*\*\*\**P* <0.0001 computed by a One-way ANOVAs with Tukey’s multiple comparisons. (**C**) Genomic position and sequences of *Nkg2d_5’E_*_Δ_ alleles (see fig. S3A legend for details). (**D**) Percentages of NKG2D+ cells in mice with the indicated *Nkg2d* genotypes, depicting one experiment each with the B1Δ and B2Δ alleles. (**E**) Comparison of percentages of % NKG2D+ NK cells (left y-axis) and mean staining intensity of NKG2D staining (normalized to +/+ mice, right y-axis) from mice with the indicated *Nkg2d* genotypes. Data are representative of two independent experiments.

**Fig. S6.**
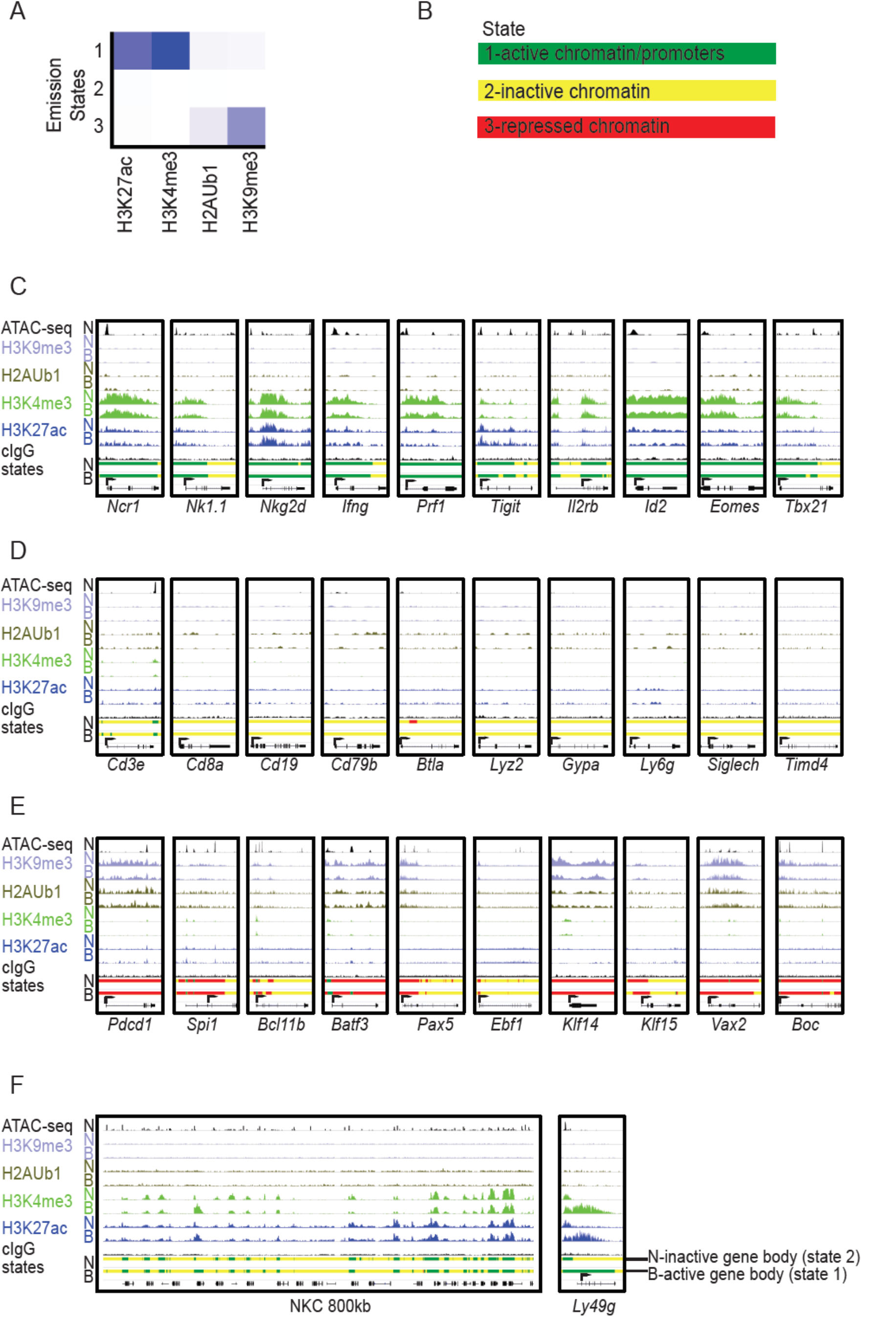
Chromatin state analysis of NK cells expressing neither (N) allele or both (B) alleles of Ly49G2. (**A**) Emission chromatin states determined by ChromHMM (*39*) in a 3 state model, based on active (H3K27ac and H3K4me3) and repressive (H2AUb1 and H3K9me3) chromatin modifications in the two NK cell populations, from CUT&RUN analyses. The vertical scale is the same for all panels A-E. (**B**) State 1 (green), defined by active modifications, is denoted “active chromatin”. State 2 (yellow) lacks both active and repressive marks and is denoted “inactive chromatin”. State 3 (red) is defined by repressive modifications and is denoted “repressed chromatin”. (**C**) IGV screenshots depicting the modifications and, at the bottom of each panel, the color-coded chromatin states of selected genes characteristically expressed by NK cells, including lineage-specific receptors, effector molecules, and transcription factors. For each modification and state, results with cells expressing neither (“N”) Ly49G2 allele, or both (“B”) are shown. (**D**) Data as in (C), except depicting selected genes encoding cell surface receptors emblematic of non-NK cell hematopoietic lineages. (**E**) Data as in (C-D), except depicting select genes expressed in non-NK cells lineages that exhibit state 3 or “repressed” chromatin either across the entire gene locus or proximal to the promoter. (**F**) Data as in (C-F), depicting the entire 800 kb segment of the NKC containing the *Ly49* and *Nkg2* loci (left) and zoomed in on the *Ly49g* locus (right). In all panels, arrows indicated the position of the annotated promoters in the EDPNew database (*33*). Vertical scales within a dataset are constant within and across panels.

**Fig. S7.**
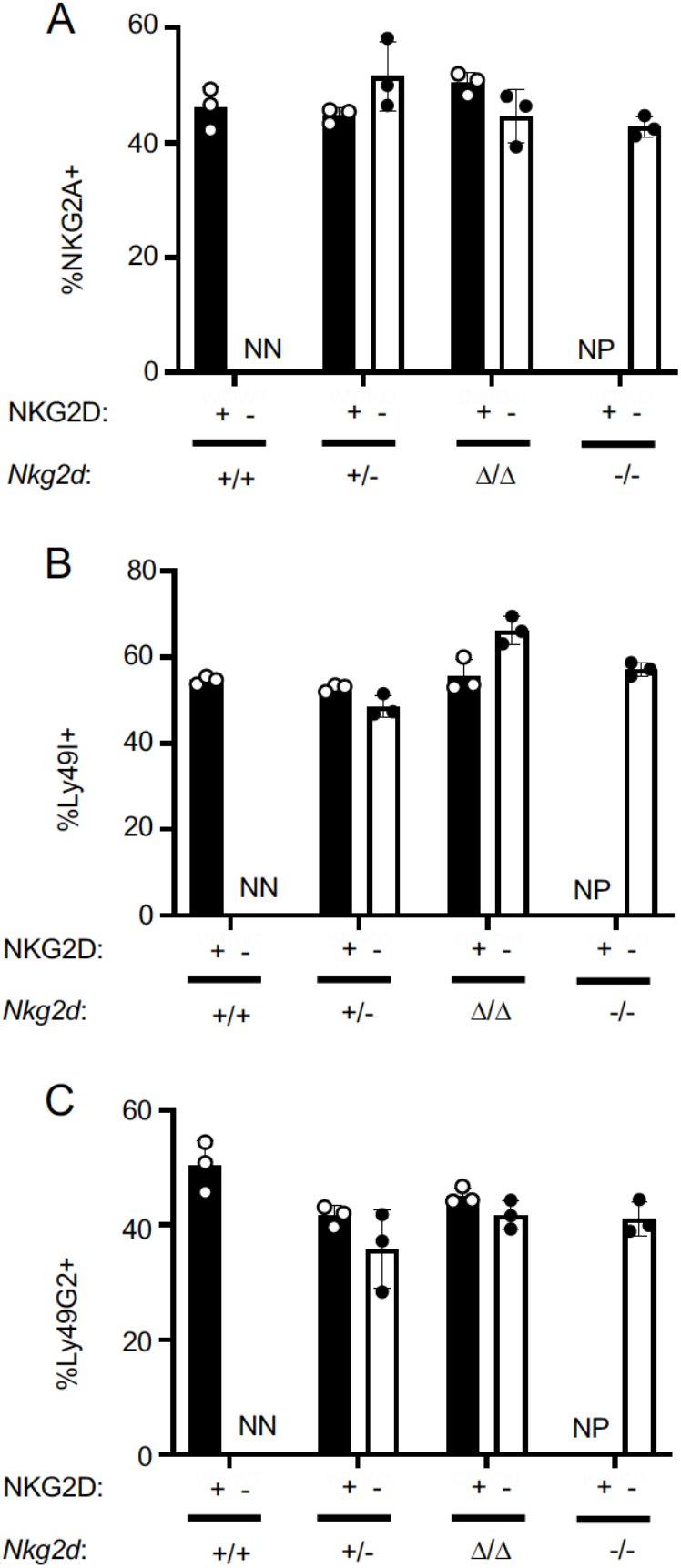
Similar patterns of NKG2A, Ly49I and Ly49G2 expression in NKG2D^+^ and NKG2D^-^ NK cells. (**A-C**) Percentages of NK cells expressing NKG2A (A), Ly49I (B) or Ly49G2 (C) among gated NKG2D^+^ and NKG2D^-^ NK cells in mice with the indicated genotypes. NN=insufficient negatives to stain; NP=no positives.

**Fig. S8.**
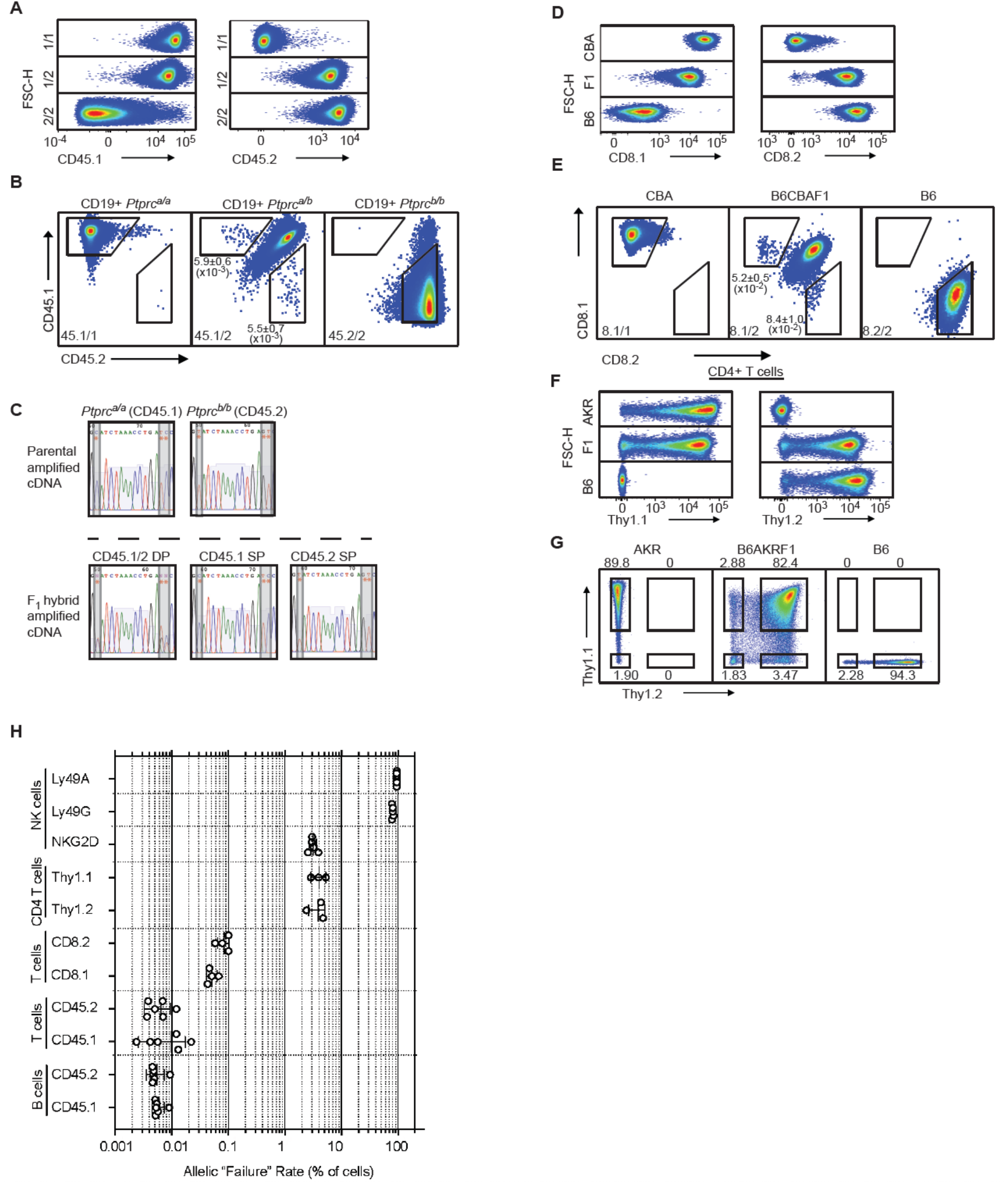
Monoallelic expression of receptors thought to be expressed by all cells in various hematopoietic lineages compared to NK cell receptors. (**A**) Scatterplot staining patterns of gated CD19+ splenic B cells from mice with the indicated genotypes with CD45.1 and CD45.2 mAbs vs FSC-H. The genotypes are as follows: 1/1: *Ptprc^a/a^*, 1/2: *Ptprc^a/b^*, and 2/2: *Ptprc^b/b^*. One mouse from each of the depicted genotypes is displayed; data are representative of 3 independent experiments. (**B**) Staining of B cells for CD45.1 and CD45.2 expression on a two-dimensional scatter plot. A single representative mouse is displayed for each depicted genotype. Gates outlining cells with monoallelic CD45 expression are shown. Mean percentages and SEMs from 3 experiments are shown in the panels. (**C**) Sanger sequence reads of amplified cDNA from *Ptprc* transcripts isolated from the indicated cells depicted in Fig. 7D. RNA was isolated from *ex vivo* expanded parental (homozygous) T-cells from *Ptprc^a/a^* and *Ptprc^b/b^* mice (top), or sorted and expanded *Ptprc^a/b^* F_1_ hybrid T-cells (bottom) expressing both alleles (CD45.1/2 DP), only the *Ptprc^a^* allele (CD45.1 SP), or only the *Ptprc^b^* allele (CD45.2 SP). Red asterisks denote the position of allele-informative SNPs between the *Ptprc^a^* and *Ptprc^b^* alleles assayed in the amplified region. (**D**) Scatterplot staining patterns of gated CD8*β*+ splenic T cells from mice with the indicated genotypes with CD8.1 or CD8.2 mAbs vs FSC-H. Data are representative of 4 independent experiments. (**E**) Two-dimensional scatter plots showing CD8.1 vs CD8.2 staining of gated CD8*β*+ cells. A single representative mouse is displayed for each genotype. Gates outlining cells with monoallelic CD8*α* expression are shown. Mean percentages and SEMs from 4 experiments from (B6 x CBA)F_1_ mice are shown in the panels. (**F**) Scatterplot staining patterns of gated CD4+ splenic T cells from B6, AKR and (B6 x AKR)F_1_ mice. Thy1.1 (left) or Thy1.2 (right) staining is depicted against FSC-H. Data are representative of 3 independent experiments. (**G**) Scatterplots of the data depicted in (F) but showing the Thy1.1 vs Thy1.2 parameters. In all cases, a single representative mouse is displayed. Data in (G) are duplicated from Fig. 7G for ease of reference. **(H)** Quantification of failure rates of selected alleles in this study. Failure rate is defined as the percentage of cells in the indicated cell population that fail to express a particular allele as measured in a genetic background that allows detection of such cells by flow cytometry. *Ly49* allelic failure rates were based on analysis in a (B6 x BALB/c)F_1_ hybrid background; *Nkg2d* alleles in mice where the opposing chromosome harbors the *Nkg2d* knockout allele; *Cd8a* alleles in (B6 x CBA)F_1_ hybrids; *Cd45* alleles in *Ptprc^a/b^* F_1_ congenic mice on the B6 genetic background; *Thy1* alleles in (B6 x AKR)F_1_ mice. Data are compiled from 3-6 mice per group from multiple experiments. The horizontal axis depicting failure rates as a percentage of cells is on a log_10_ scale. In each case, error bars represent the SEM.

**Table S1.**
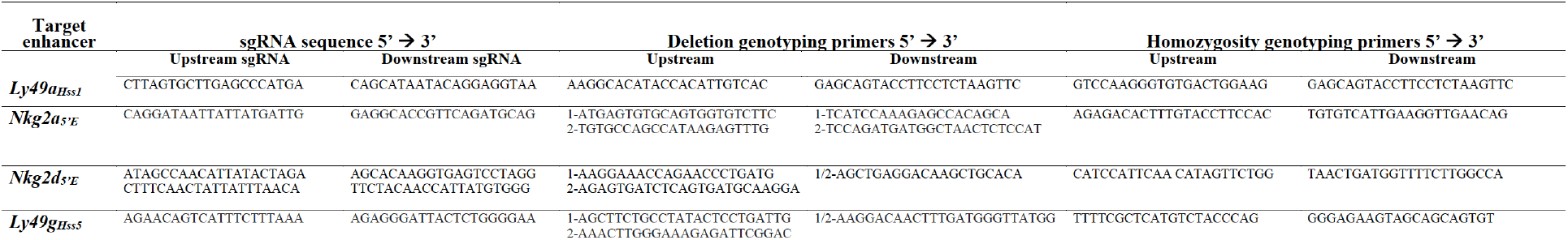
Guides and primers used to generate and genotype CRISPR/Cas9-edited mice. Guide RNAs (sgRNAs) used to generate germline enhancer deletion mice via electroporation or microinjection are displayed. A flanking guide pair was used to delete the indicated enhancer, except for in the case of *Nkg2d_5’E_*, where two sets of flanking guides were used (all four sgRNAs were simultaneously delivered to embryos). Primers used to genotype mice carrying a deletion allele and mice lacking a WT allele are also shown. These primers allow delineation of WT, heterozygous and homozygous enhancer deletion animals with respect to the indicated enhancer element. More than one primer is shown if PCR was performed as a nested reaction; “1” indicates use in the first amplification and “2” indicates use in the subsequent amplification.

**Table S2.**
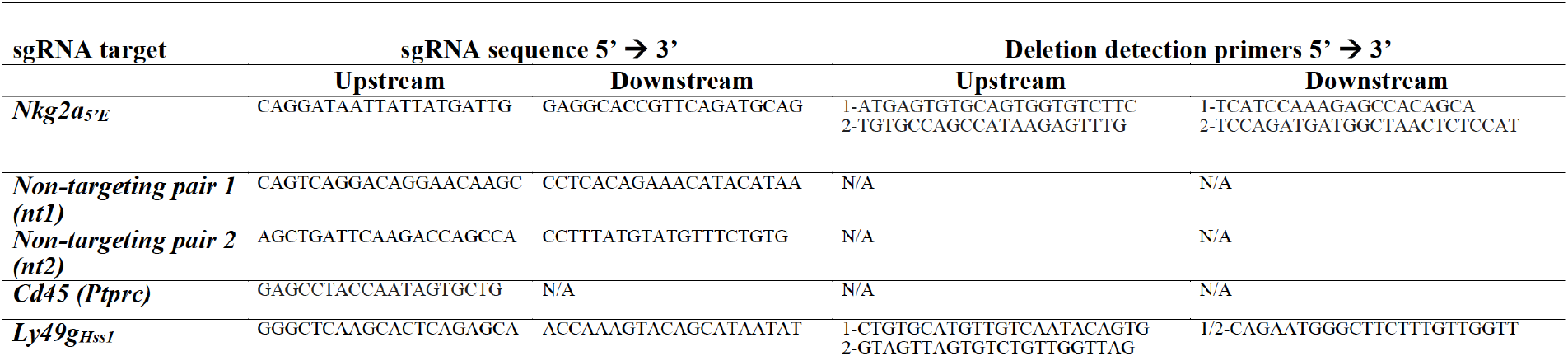
Guides and primers used for ex vivo NK cell editing. Guide RNAs (sgRNAs) used in the *ex vivo* NK cell enhancer deletion assay are displayed. Non-targeting sgRNA pairs 1 (nt1) and 2 (nt2) were used as negative controls in fig. S3A. Primers used to detect the presence of the intended deletion in nucleofected NK cells using the indicated sgRNAs used are also shown. More than one primer is shown if PCR was performed as a nested reaction; “1” indicates use in the first amplification and “2” indicates use in the second amplification.

**Table S3.**
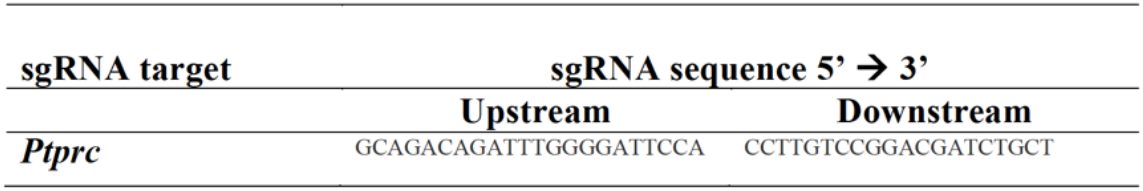
Primers used to amplify *Ptprc* PCR products. Intron-spanning primers detecting a region of the *Ptprc* transcript containing 3 allele-informative SNPs.

